# Skeletal cell YAP and TAZ redundantly promote bone development by regulation of collagen I expression and organization

**DOI:** 10.1101/143982

**Authors:** Christopher D. Kegelman, Devon E. Mason, James H. Dawahare, Genevieve D. Vigil, Scott S. Howard, Teresita M. Bellido, Alexander G. Robling, Joel D. Boerckel

## Abstract

The functions of the transcriptional co-activators YAP and TAZ in bone are controversial. Each has been observed to either promote or inhibit osteogenesis *in vitro*, while their roles in bone development are unknown. Here we report that combinatorial YAP/TAZ deletion from skeletal cells in mice caused osteogenesis imperfecta with severity dependent on targeted cell lineage and allele dosage. Osteocyte-conditional deletion impaired bone accrual and matrix collagen, while allele dosage-dependent deletion from all osteogenic lineage cells caused spontaneous fractures, with neonatal lethality only in dual homozygous knockouts. We identified putative target genes whose mutation in humans causes osteogenesis imperfecta and which contain promoter-proximate binding domains for the YAP/TAZ co-effector, TEAD4. Two candidates, Col1a1 and SerpinH1, exhibited reduced expression upon either YAP/TAZ deletion or YAP/TAZ-TEAD inhibition by verteporfin. Together, these data demonstrate that YAP and TAZ redundantly promote bone matrix development and implicate YAP/TAZ-mediated transcriptional regulation of collagen in osteogenesis imperfecta.

## INTRODUCTION

Bone is a living hierarchical composite whose form and function depend not only on the composition of the matrix, but also its microstructure, which are controlled during development by skeletal cell lineage progression and by osteocyte-coordinated matrix deposition and remodeling. Various genetic, hormonal, or environmental abnormalities can impair these processes, leading to debilitating diseases including osteoporosis and osteogenesis imperfecta. However, the molecular mechanisms governing cell fate and matrix production in bone remain poorly understood, limiting therapeutic intervention. Several transcriptional programs have been described as essential regulators of bone development, but current understanding is insufficient to fully explain the heterogeneity found in congenital and acquired bone diseases^1–3^. In this study, we sought to define the function of the Hippo pathway effectors YAP and TAZ in bone development.

### YAP/TAZ functional diversity

Yes-associated protein (YAP) and Transcriptional co-activator with PDZ-binding motif (TAZ; also known as WWTR1) are paralogous transcriptional co-activators that display either equivalent or divergent functions, depending on cell type and context^4^. While they possess transcription activation domains, they lack DNA-binding domains and require interaction with co-factors for transcriptional activity^5^. Their most well-studied interactions are with the transcriptional enhancer activator-domain (TEAD) family proteins, which themselves lack activation domains, providing specificity for YAP/TAZ-TEAD signaling^6^. Many other co-effectors are known, including Runx2^7^, β-catenin^8–10^, and Smad2/3^11,12^, each of which is known to contribute to bone development and osteoprogenitor lineage progression^13–17^ Thus, independent pathways that regulate coincident activation of these various binding partners could provide additional layers of contextual specificity in bone. Further, as paralogs, the YAP and TAZ proteins also possess structural differences (reviewed in Ref.^18^) that enable distinct protein interactions to confer unique physiological functions of YAP vs. TAZ. Notably, global YAP deletion in mice is embryonic lethal (E8.5) due to impaired yolk sac vasculogenesis^19^, while the global TAZ knockout lives to maturity with modest skeletal defects^20^ demonstrating conclusive gene-specific functions. However, in other contexts, they exhibit clear functional homology, with either protein capable of compensation for the other^21,22^.

### YAP and TAZ function in bone: conflicting evidence

Roles for YAP and TAZ in osteogenesis were first described in 2004 and 2005, respectively^23,24^. YAP was reported to suppress osteoblastic differentiation through sequestration and transcriptional repression of Runx2^23^, while TAZ was identified as a Runx2 co-activator and an inhibitor of the adipogenic nuclear receptor, PPARy^24,25^. A subsequent study found that overexpression of a constitutively-active YAP mutant in marrow stromal cells (MSCs) promoted osteogenic differentiation even under conditions more favorable for adipogenesis^26^. In contrast, another report identified that YAP overexpression inhibited osteogenesis in MSCs by suppressing activation of WNT target genes^27^. The role of TAZ in osteogenic differentiation appears similarly complicated with a recent report demonstrating that overexpression of active-mutant TAZ inhibited canonical WNT/β-catenin signaling^28^, while another suggested that TAZ promoted osteogenic differentiation through the canonical WNT pathway^29^. *In vivo*, osteoblast-specific overexpression of TAZ promoted bone formation with higher expression levels of Runx2^30^, while YAP overexpression in chondrocytes impaired endochondral bone formation^31^. These data suggest the importance of these proteins in osteogenic differentiation and mineralized matrix production, but their combinatorial physiological roles in bone development remain unknown.

## RESULTS

### YAP/TAZ expression and deletion in bone

To determine YAP/TAZ expression profiles in bone, we immunostained YAP and TAZ in the growth plate and cancellous and cortical bone of 8 week-old C57Bl6/J mouse femora. YAP and TAZ immunolocalized in hypertrophic chondrocytes, osteoblasts, and osteocytes with minimal detectable expression in quiescent or proliferating chondrocytes (Fig. S1A). Based on these expression patterns, we chose to assess the physiological roles of YAP and TAZ by combinatorial conditional ablation^22^ at two stages in the skeletal cell sequence: 1) all cells of the skeletal lineage using Osterix1-GFP::Cre (Osx1-Cre)^32^ and 2) osteocytes using 8kb-DMP1-Cre (Dmp1-Cre)^33,34^. We selected a breeding strategy that yielded littermates with variable YAP/TAZ allele dosage (Table S1). To verify Cre-mediated recombination and deletion of YAP and TAZ, we assessed mRNA expression in femoral bone preparations by qPCR (Fig. S1D-F), and verified the absence of expression in bone cells in conditional knockout mice by IHC. YAP/TAZ expression was reduced by 50-80% in bone preparations from homozygous knockout mice with either Osx1-Cre- or DMP1-Cre-mediated excision (Fig. S1B-G).

**Table S1:** Genotypes evaluated

| Osx1-GFP::Cre | YAP/TAZ alleles | 8kb-DMP1-Cre | YAP/TAZ alleles |
| --- | --- | --- | --- |
| Yap<sup>fl/fl</sup>;Taz<sup>fl/fl</sup> | YAP<sup>WT</sup>;TAZ<sup>WT</sup> | Yap<sup>fl/fl</sup>;Taz<sup>fl/fl</sup> | YAP<sup>WT</sup>;TAZ<sup>WT</sup> |
| Yap<sup>fl/+</sup>;Taz<sup>fl/+</sup>;Osx1<sup>cre/+</sup> | YAP<sup>CHET</sup>;TAZ<sup>CHET</sup> | Yap<sup>fl/+</sup>;Taz<sup>fl/+</sup>;DMP1<sup>cre/+</sup> | YAP<sup>CHET</sup>;TAZ<sup>CHET</sup> |
| Yap<sup>fl/fl</sup>;Taz<sup>fl/+</sup>;Osx1<sup>cre/+</sup> | YAP<sup>CKO</sup>;TAZ<sup>CHET</sup> | Yap<sup>fl/fl</sup>;Taz<sup>fl/+</sup>;DMP1<sup>cre/+</sup> | YAP<sup>CKO</sup>;TAZ<sup>CHET</sup> |
| Yap<sup>fl/+</sup>;Taz<sup>fl/fl</sup>;Osx1<sup>cre/+</sup> | YAP<sup>CHET</sup>;TAZ<sup>CKO</sup> | Yap<sup>fl/+</sup>;Taz<sup>fl/fl</sup>;DMP1<sup>cre/+</sup> | YAP<sup>CHET</sup>;TAZ<sup>CKO</sup> |
| Yap<sup>fl/fl</sup>;Taz<sup>fl/fl</sup>;Osx1<sup>cre/+</sup> | YAP<sup>CKO</sup>;TAZ<sup>CKO</sup> | Yap<sup>fl/fl</sup>;Taz<sup>fl/fl</sup>;DMP1<sup>cre/+</sup> | YAP<sup>CKO</sup>;TAZ<sup>CKO</sup> |

**Supplementary Figure 1.**
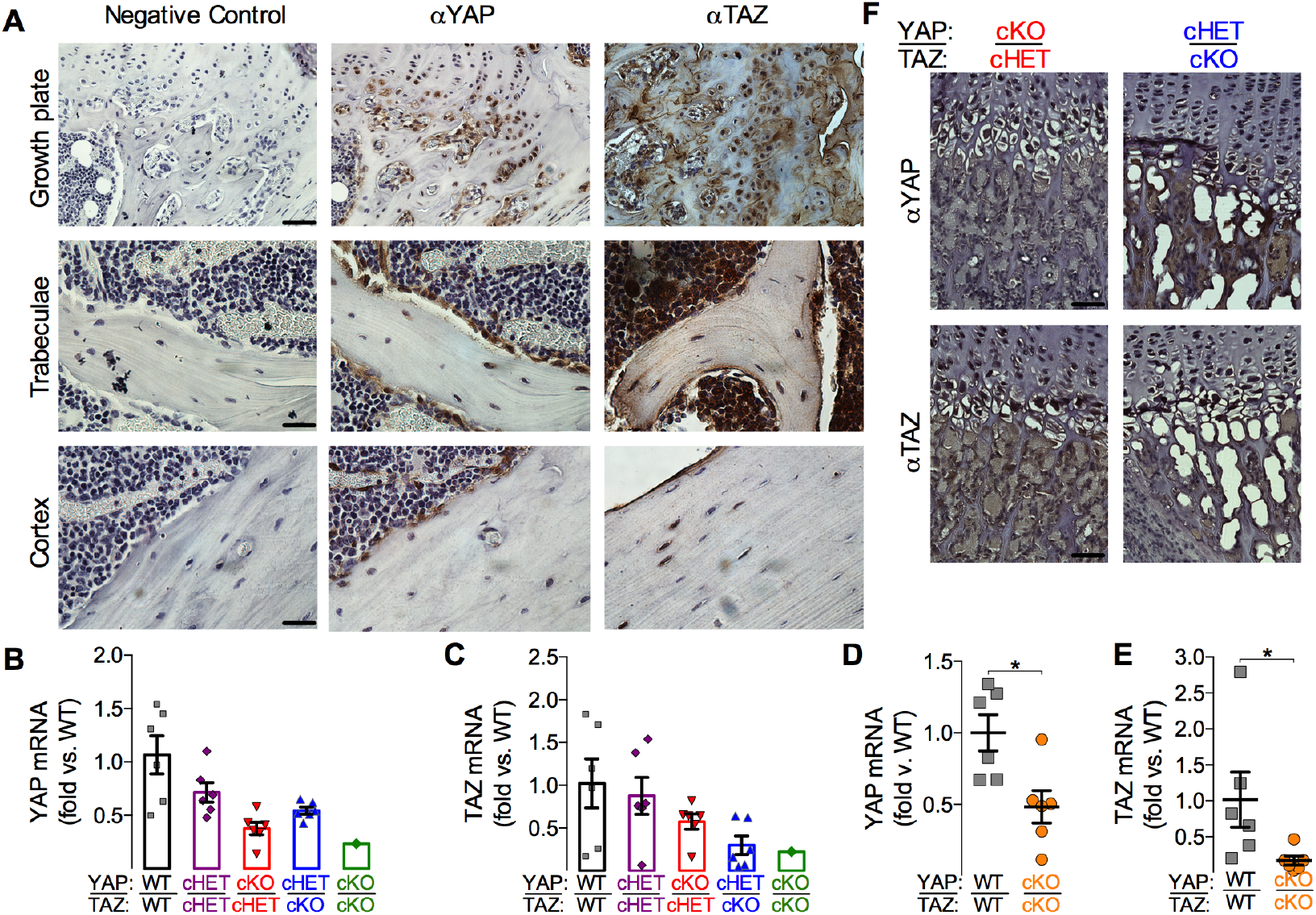
Immunolocalization of YAP/TAZ expression and YAP/TAZ recombination efficiency. *In vivo* YAP/TAZ expression and knockdown efficiency were assessed by immunohistochemistry (IHC) and quantitative (qPCR). **A**) YAP/TAZ were expressed in hypertrophic chondrocytes, osteoblasts, and osteocytes in bone. Scale bars in the first column are 50 microns for the growth plate, and 100 microns for magnified images of trabecular and cortical bone. YAP (**B,D**) and TAZ (**C,E**) transcript expression in Osterix- and DMP1-conditional knockout mouse bone. (**F**) IHC verification of YAP and TAZ knockdown at the protein level in Osterix-conditional cKO bone. Scale bars in the first column are 50 microns.

### Osterix-conditional YAP/TAZ cKO: neonatal lethality and hypermineralization

All osterix-conditional knockouts and littermate controls were born at expected Mendelian ratios, but dual homozygous conditional deletion (YAP^cKO^;TAZ^cKO^) caused neonatal asphyxiation secondary to ribcage malformation and fracture (Fig. 1A-C), resulting in 75% mortality at postnatal day 0 (P0) and 99% by P7 (Fig. 1B). Only one female YAP^cKO^;TAZ^cKO^ mouse lived to P56 for each endpoint analysis. YAP^cKO^;TAZ^cKO^ neonates exhibited spinal scoliosis, cranial vault deformity, and spontaneous fractures of the ribs, tibia, femur, radius and ulna (Fig. 1A,C-E). Spontaneous extremity fractures were not present in other genotypes at P0 (Fig. 1A). Littermate neonates displayed a trend of reduced whole-skeleton bone volume (Fig. 1B; p=0.05, ANOVA) and significantly elevated bone tissue mineral density (Fig. 1C; p<0.01, ANOVA) with decreasing YAP/TAZ allele dosage. Osterix-conditional YAP/TAZ deletion also significantly reduced birth weight and intact femoral length in an allele dosage-dependent manner (Fig. S2A). Males and females exhibited similar phenotypes (Fig. S2B).

**Figure 1.**
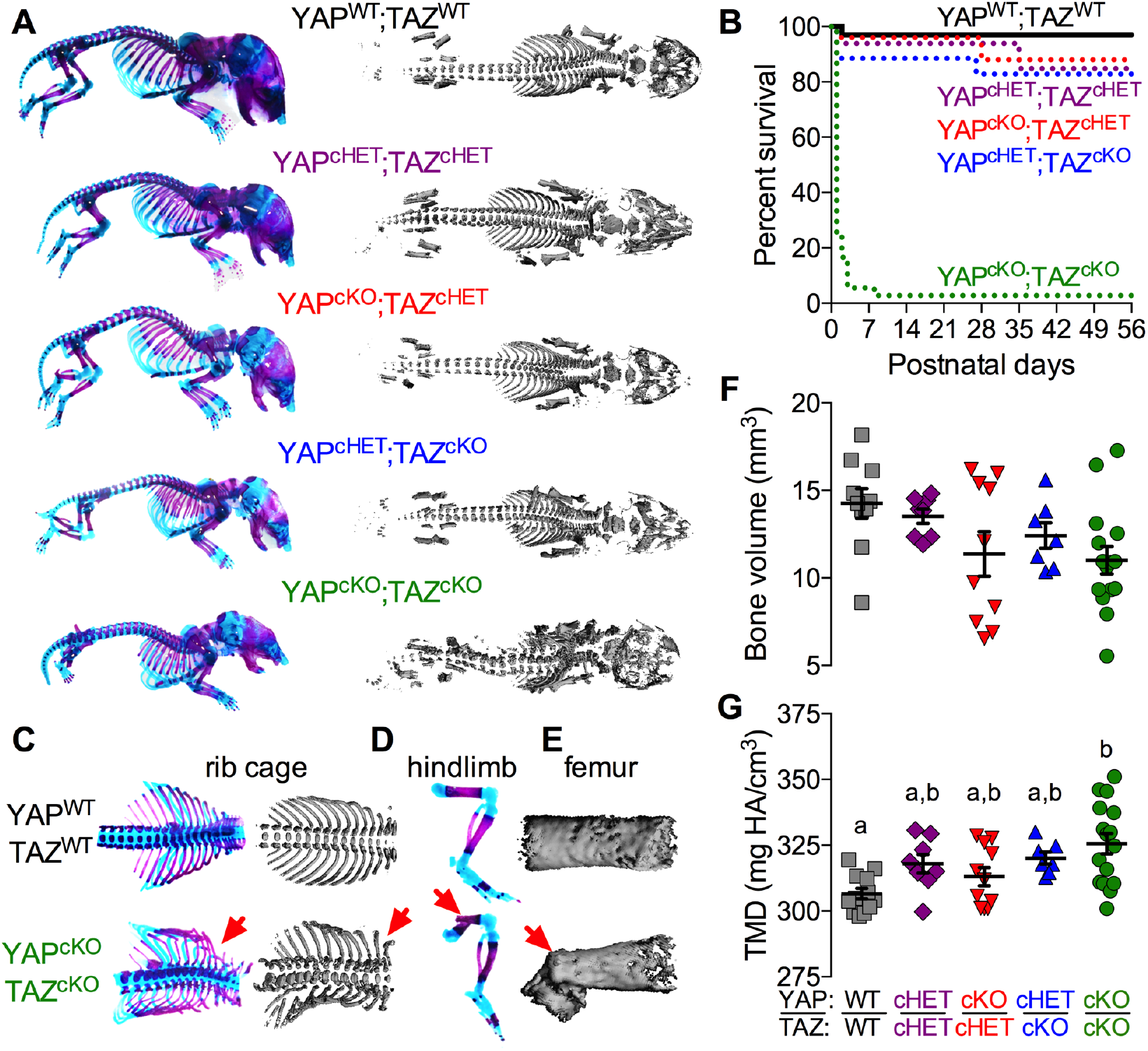
Combinatorial YAP/TAZ ablation from Osterix-expressing cells caused allele dosage-dependent perinatal skeletal deformity and lethality. Skeletal structures of littermate mice were evaluated at postnatal day 0 (P0). **A**) Whole body skeletal preparations of osterix-conditional YAP/TAZ knockouts and controls stained with Alcian blue/Alizarin red and microCT reconstructions reveal progressive skeletal malformation with decreasing allele dosage. **B**) Survival curves for each genotype show 99% lethality of YAP^cKO^;TAZ^cKO^ mice by P56. **C-E**) Skeletal preparations and microCT reconstructions of rib cages, hindlimbs, and femora, respectively, illustrate spontaneous perinatal fractures in YAP^cKO^;TAZ^cKO^ mice. **F**) P0 whole skeleton bone volume was not significantly altered by YAP/TAZ deletion. **G**) P0 whole skeleton tissue mineral density (TMD) increased with YAP/TAZ allele deletion. Data are presented as individual samples with lines corresponding to the mean and standard error of the mean (SEM). Sample sizes, n = 7-15.

**Supplementary Figure 2.**
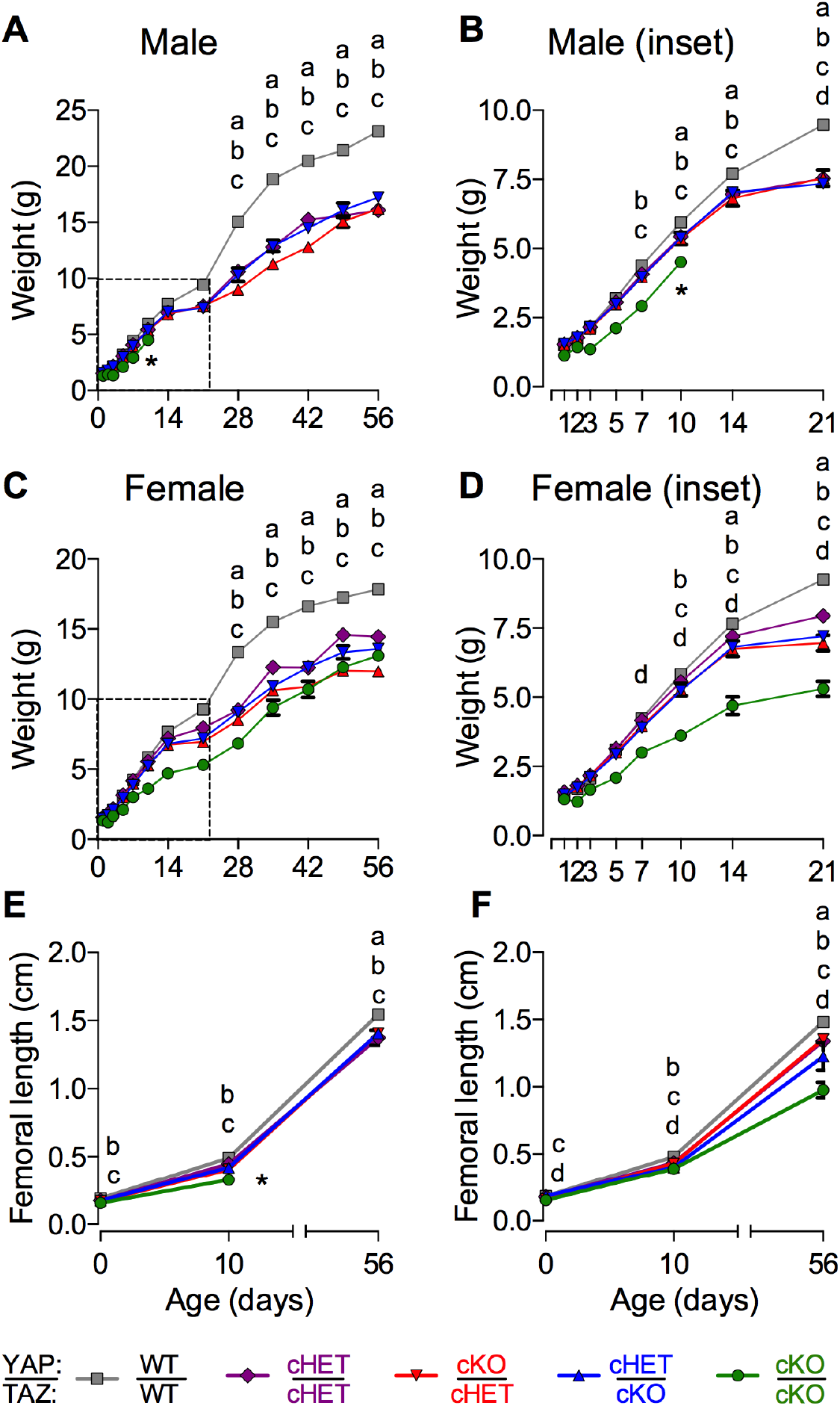
Allele-dose dependent YAP/TAZ ablation from Osterix-expressing cells in 8-week-old mice reduced overall and long bone growth independent of sex. Allele-dosage-dependent YAP/TAZ ablation from Osterix-expressing cells reduced body weight in male (**A, B**) and female (**C,D**) mice and reduced femoral length (**E, F**). Sample sizes, n = 8 for both genders and genotypes except n = 1 for YAP^cKO^;TAZ^cKO^ male and n = 2 for YAP^cKO^;TAZ^cKO^ females. * indicates death of male YAP^cKO^;TAZ^cKO^ mouse at P10.

### Osterix-conditional YAP/TAZ cKO: spontaneous neonatal long bone fractures & defective endochondral bone formation

A single copy of either gene rescued neonatal lethality, with 83 and 85% of YAP^cHET^;TAZ^cKO^ and YAP^cKO^;TAZ^cHET^ mice surviving to terminal analysis at P56, respectively. However, between P1 and P10, both YAP^cHET^;TAZ^cKO^ and YAP^cKO^;TAZ^cHET^ mice sustained spontaneous femoral and other bone fractures (Fig. 2A-B), with significantly increased femoral fracture incidence in the YAP^cKO^;TAZ^cHET^ mice (p<0.01, χ^2^ test; Fig. 2C). Fractures healed by endochondral repair in all groups, though YAP^cHET^;TAZ^cKO^ and YAP^cKO^;TAZ^cKO^ calluses exhibited numerous empty lacunae in the hypertrophic transition zone, suggesting increased hypertrophic chondrocyte death or insufficient progenitor cell recruitment (Fig. 2C,D). Consistently, staining of Osterix-positive cells was qualitatively reduced in the transition zone of YAP^cKO^;TAZ^cKO^ growth plates, but we found no significant differences in growth plate morphology at P10 (Fig. 2E,F). Histomorphometric analysis of P56 growth plates did not reveal significant differences in proliferative or hypertrophic zone thickness, though the single YAP^cKO^;TAZ^cKO^ mouse that survived to P56 exhibited a lower relative hypertrophic zone thickness (Fig. 2G,H). Osterix-conditional YAP/TAZ deletion did not alter osteocyte density (Ot.N/B.Ar) at P56 (Fig. 2I,J).

**Figure 2.**
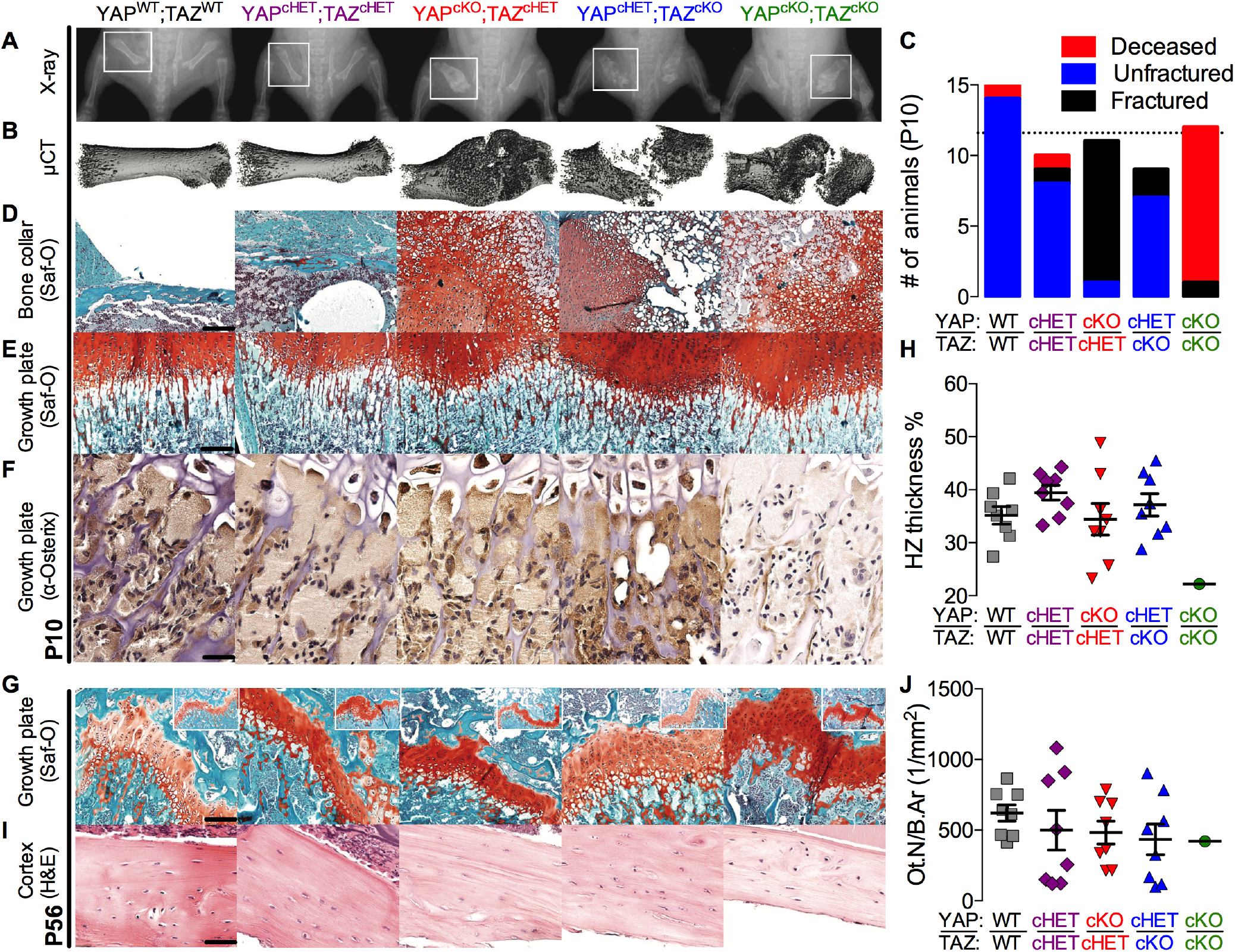
YAP/TAZ ablation from Osterix-expressing cells induced spontaneous neonatal femoral fractures and impaired endochondral bone formation. Representative radiographs (**A**) with matched microCT reconstructions of femoral fracture calluses at P10 (**B**). **C**) Quantification of the number of femoral fractures demonstrated significantly increased fracture incidence in YAP^cko^;TAZ^cHET^ mice. **D**) Safranin-O/fast green (Saf-O) staining of mid-diaphysis bone collar or fracture callus and (**E**) growth plates of matched P10 femora. Scale bar = 50 μm. **F**) Representative micrographs of P10 distal femur growth plates immunostained for osterix-positive cells (brown). Scale bar = 25 μm. **G**) Representative Safranin-O/fast green-stained distal femoral growth plates at P56. Scale bar = 50 μm. Inset: Saf-O/fast green (Inset width = 1.5 mm). **H)** Histomorphometric quantification of P56 hypertrophic zone thickness as percentage of total growth plate thickness (HZ thickness %). **I)** Representative micrographs of P56 mid-diaphyseal cortical bone stained by H&E. **J)** Histomorphometric quantification of osteocyte number per bone area (Ot.N/B.Ar). Data presented as individual samples with lines corresponding to the mean and standard error of the mean (SEM). Sample sizes, n = 3-6 at P10, n = 8 at P56, except YAP^cKO^;TAZ^cKO^ n = 1.

### Osterix-conditional YAP/TAZ cKO: reduced cortical and cancellous microarchitectural properties

YAP/TAZ deletion from all cells of the skeletal lineage reduced cancellous (Fig. 3A,B) and cortical bone (Fig. 3C,D) volume and shape, according to allele dosage. Distal femur metaphyseal cancellous bone exhibited reduced trabecular bone volume fraction (BV/TV), thickness (Tb.Th), and number, and increased spacing and structural model index (SMI, indicative of more rod-like trabeculae) (Fig. 3B, Fig. S3A). The cumulative distribution of trabecular thicknesses shifted in an allele dosage-dependent manner, indicating reduced numbers of both small and large trabeculae (Fig. 3B, S3B). Volumetric bone mineral density was not altered, suggesting an increase in local tissue mineral density proportional to the decrease in trabecular bone volume (Fig. 3B). Mid-diaphyseal femoral cortical bone (Fig. 3C,D) similarly exhibited reduced thickness, area, and moment of inertia (I) in cKO mice, attributable primarily to reduced periosteal bone accumulation as indicated by moderate reductions in medullary area (Me.Ar) compared to cortical bone area (B.Ar) and total area (T.Ar). Consistent with the observations of vBMD in the cancellous compartment, cortical tissue mineral density (Ct.TMD) was significantly increased in an allele-dosage dependent manner; however, unlike the cancellous bone, the increase in tissue mineral density of the cortical bone was insufficient to normalize bone mass lost by reduced bone volume. Together, these data demonstrate that both YAP and TAZ contribute positively to bone quantity and microarchitecture in both cancellous and cortical compartments.

**Figure 3.**
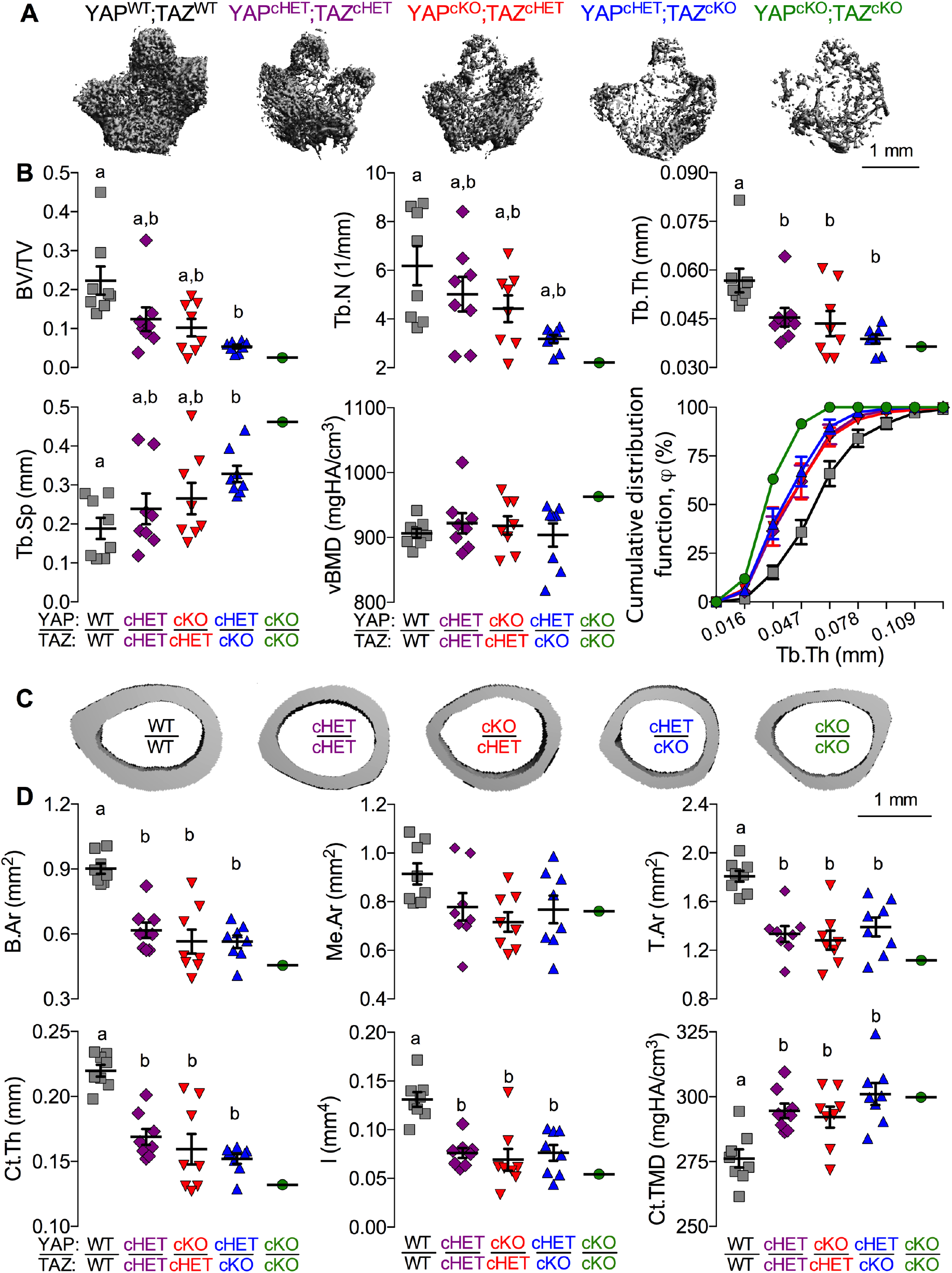
YAP/TAZ ablation from Osterix-expressing cells altered bone microarchitectural properties in a manner dependent on allele dosage. Femora from 8 week-old osterix-conditional YAP/TAZ littermates were evaluated by microCT analysis. **A)** Representative microCT reconstructions of metaphyseal cancellous bone, arranged in decreasing allele dosage. **B)** Cancellous bone microarchitectural parameters were impaired according to YAP/TAZ allele dosage: bone volume fraction (BV/TV), trabecular thickness (Tb.Th), number (Tb.N) and spacing (Tb.Sp), and volumetric bone mineral density (vBMD). The cumulative distribution function, *ϕ*, demonstrated a reduced percentage of trabeculae smaller than a given thickness (on the abscissa) with increasing allele deletion. **C)** Representative microCT reconstructions of the mid-diaphyseal cortex, arranged in decreasing allele dosage. **D)** Cortical cross-sectional properties were reduced in cKO mice: bone area (B.Ar), medullary area (Me.Ar), total area (T.Ar), cortical thickness (Ct.Th), moment of inertia in the direction of bending (I), and cortical tissue mineral density (Ct.TMD). Data are presented as individual samples with lines corresponding to the mean and standard error of the mean (SEM). Sample sizes, n=8 except YAP^cKO^;TAZ^cKO^ n = 1. Scale bars indicate 1 mm for microCT reconstructions.

**Supplementary Figure 3.**
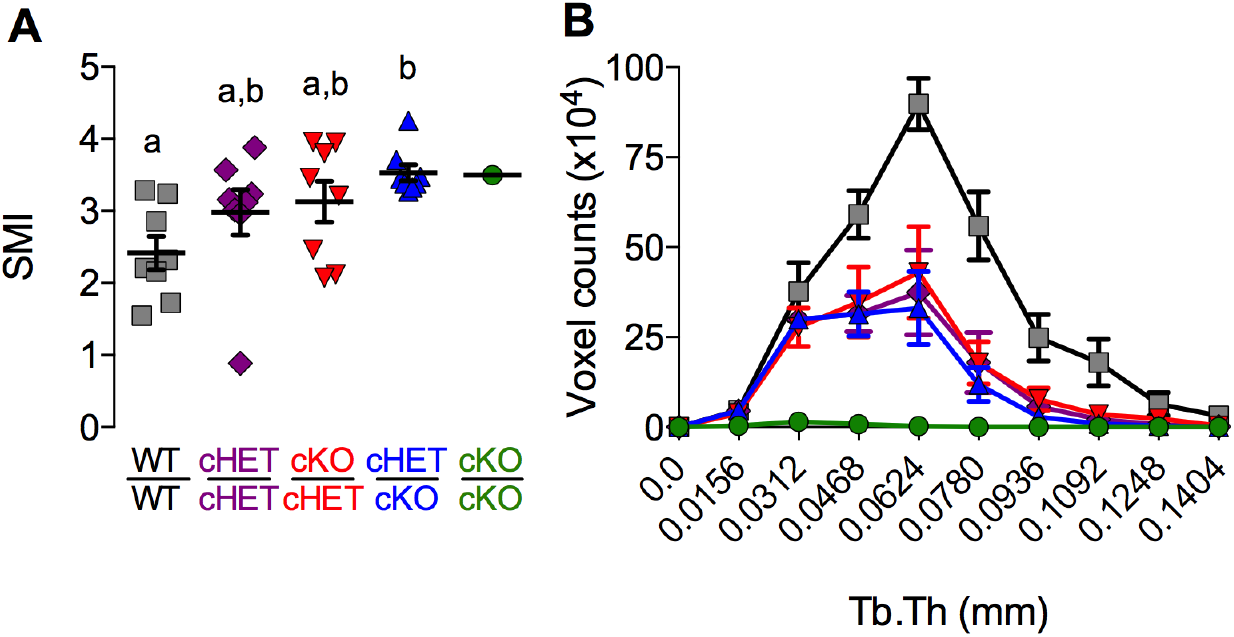
Allele-dose dependent YAP/TAZ ablation from Osterix-expressing cells in 8-week-old mice altered trabecular architecture. **A)** Structural model index (SMI) **B)** Trabecular thickness (Tb.Th) histogram distributions. Sample sizes, n= 8 except YAP^cKO^;TAZ^cKO^ n = 1.

### Osterix-conditional YAP/TAZ cKO: reduced intrinsic bone mechanical properties and matrix collagen content and microstructure

To determine whether Osterix-conditional YAP/TAZ deletion impaired adolescent bone matrix quality, we performed three-point bend testing to failure (Fig. 4A,B) on each femur previously analyzed by microCT. The extrinsic bone properties (i.e., maximum load at failure, bending stiffness, work to max load, and work to failure; Fig. 4C-F, respectively) depend on both the intrinsic mechanical properties of the bone matrix and the bone amount and cross-sectional distribution. Since the assumptions of Euler-Bernoulli beam theory are decidedly not met in three-point bend testing of mouse long bones^35,36^, we performed an analysis of covariance (ANCOVA) using linear regression (Fig. 4G,H) to decouple the contributions of bone quantity and distribution from the mechanical behavior^37^. If the variability in extrinsic mechanical properties is best predicted by individual regression lines for each genotype, this would indicate differences in intrinsic matrix mechanical properties between genotypes; however, a best-fit by a single regression line for all groups would indicate that differences in extrinsic behavior are sufficiently described merely by changes in bone geometry^37^. We found that individual regression lines for each genotype best predicted maximum load at failure, indicating significant differences in intrinsic failure properties (Fig. 4G). In contrast, a single regression line best fit the stiffness data (Fig. 4H), indicating that the differences in stiffness can be attributed to changes in moment of inertia rather than intrinsic matrix elastic properties.

**Figure 4.**
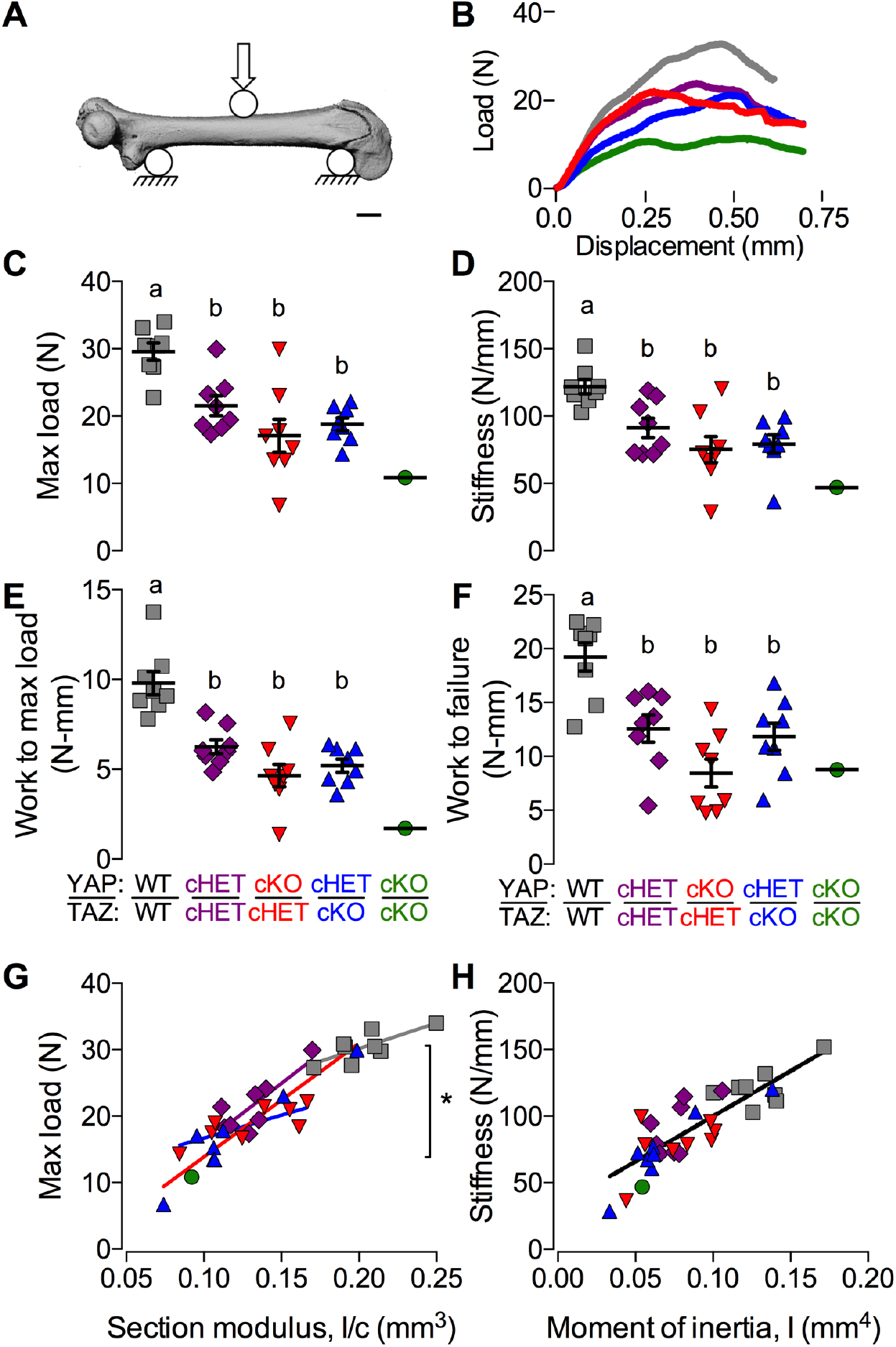
YAP/TAZ ablation from Osterix-expressing cells reduced intrinsic bone failure properties. Femora from 8 week-old osterix-conditional YAP/TAZ littermates were tested in three point bending (**A**) to failure. **B)** Representative load-displacement curves collected during testing. **C-F)** YAP/TAZ deletion reduced extrinsic mechanical properties measured from the load-displacement curves including maximum load (**C**), stiffness (**D**), work to maximum load (**E**) and work to failure (**F**). ANCOVA analysis accounting for bone geometry revealed significant differences in intrinsic failure properties (**G**), but not intrinsic elastic properties (**H**). Data are presented as individual samples with lines corresponding to the mean and standard error of the mean (SEM). Sample sizes, n = 8 except YAP^cKO^;TAZ^cKO^ n = 1.

As a composite material, bone mechanical behavior is determined predominantly by its two primary matrix components: mineral and collagen. By microCT scanning prior to mechanical testing, we noted above that femora from mice with Osterix-conditional YAP/TAZ deletion exhibited moderate hypermineralization (Fig. 3D). Next, to characterize the bone matrix collagen in these same samples, we performed polarized light microscopy and second harmonic imaging microscopy (SHIM)^38^. Both polarized-light microscopy of Picrosirius red-stained sections (Fig. 5A) and second harmonic imaging (Fig. 5B) demonstrated that YAP/TAZ deletion significantly reduced collagen content and organization (Fig. 5C), and produced a trend of reduced fiber directionality (Fig. S3A,B). Therefore, to determine the contributions of geometry, mineralization, and collagen content and microstructure to bone mechanical behavior, we performed a best-subsets correlation analysis to identify significant predictors based on Akaike’s information criterion (AIC)^39,40^. For both elastic (Fig. 5D) and failure (Fig. 5E) properties, bone tissue mineral density was not a significant predictor; however, moment of inertia and SHG intensity significantly improved model capability to explain variation (R_adj_^2^ = 73 and 88% for stiffness and max load, respectively) and reduced AIC (Fig. S4A,C). Addition of TMD did not improve predictive power or model quality (Fig. S4B,D).

**Figure 5.**
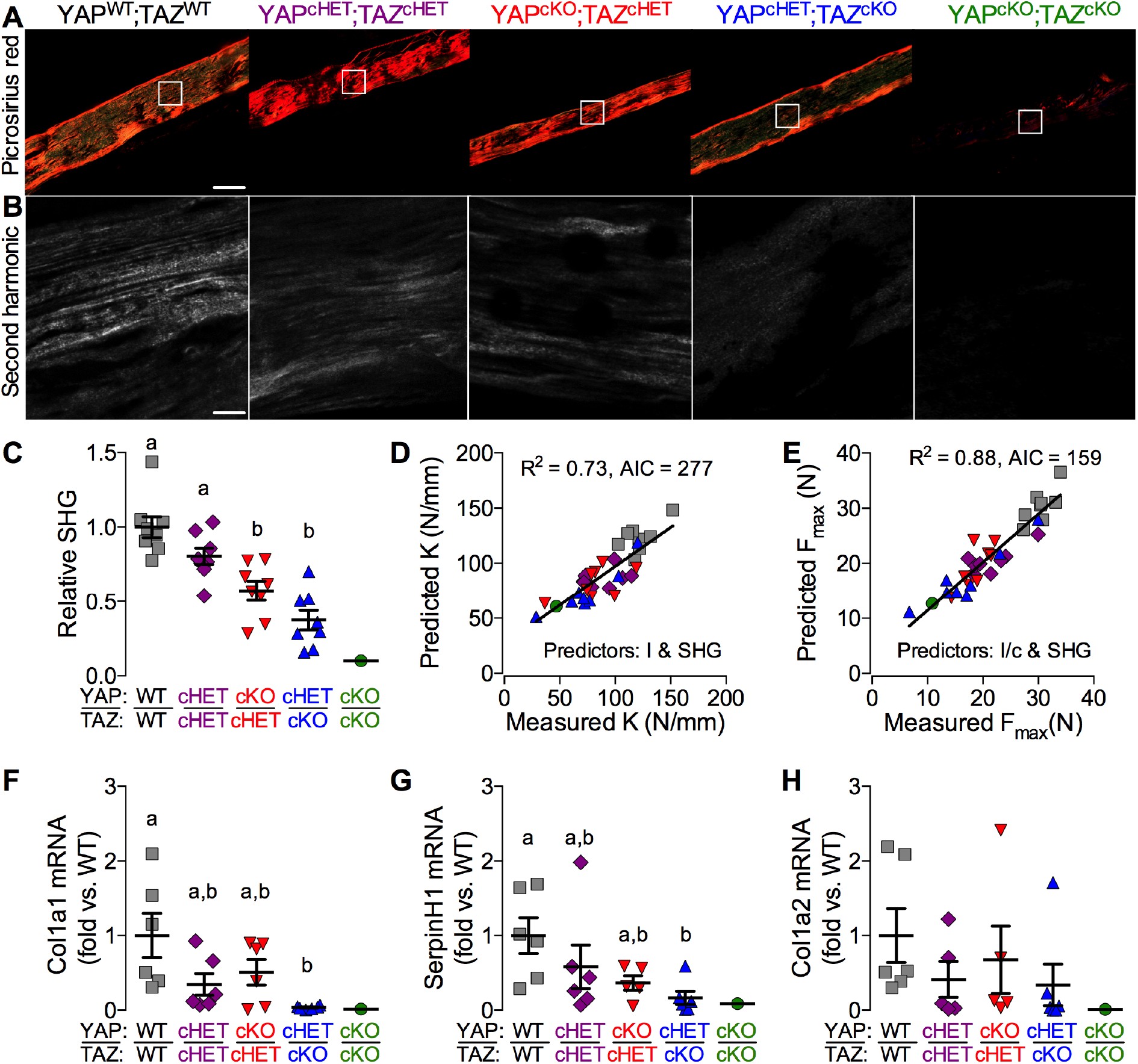
YAP/TAZ ablation from Osterix-expressing cells reduced bone matrix collagen content, organization, and gene expression. Imaging of matrix collagen and quantification of collagen-associated gene expression were performed on two sets of femora from 8 week-old osterix-conditional YAP/TAZ littermates. **A,B**) Representative polarized light (**A**) and second harmonic generation (SHG; **B**) microscopy images from cortical bone tissue sections of three-point bend tested femora. **C)** Second harmonic generation (SHG) intensity, relative to WT, was reduced according to allele-dosage. **D,E)** Best subsets regression analyses indicating significant contributions of both bone geometry and collagen content and microstructure, but not tissue mineral density, to both elastic (**D**) and failure (**E**) mechanical properties. **F-H)** Quantitative PCR amplification of genes identified as bone matrix collagen regulators with putative YAP/TAZ co-factor binding domains. Osterix-conditional YAP/TAZ deletion reduced mRNA expression of both Col1a1 (**F**) and SerpinH1 (**G**), but not Col1a2 (**H**). Sample sizes, n = 8 and 6 for SHG and qPCR analyses, respectively; except in both cases YAP^cKO^;TAZ^cKO^ n = 1. Scale bars indicate 100 *μ* m and 25 *μ* m in Picrosirius red and SHG images, respectively.

**Supplementary Figure 3.**
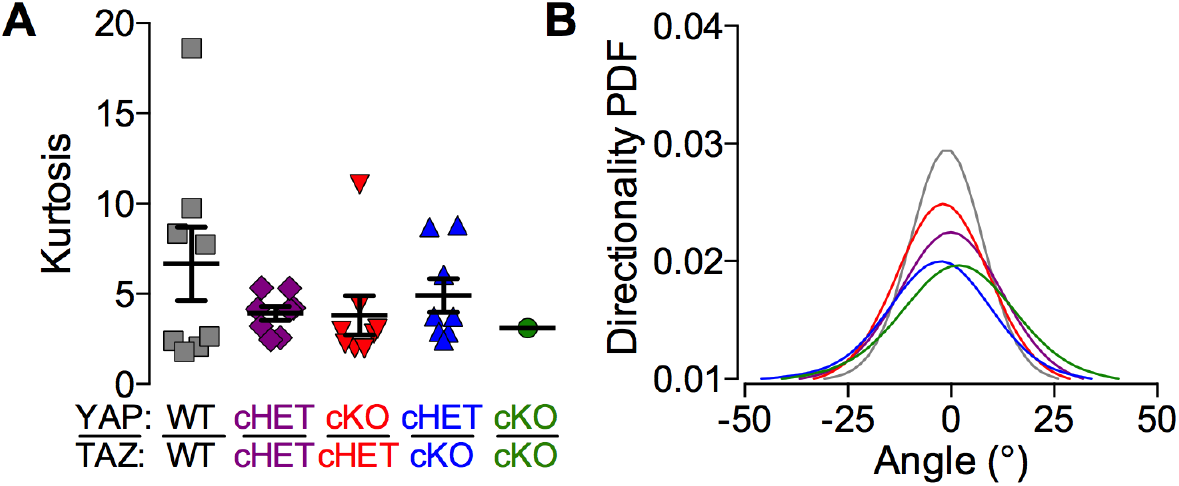
Allele-dose dependent YAP/TAZ ablation from Osterix-expressing cells in 8-week-old mice did not significantly alter collagen fiber directionality. (**A**) Kurtosis quantification of fiber directionality distributions with (**B**) accompanying mean directionality distribution probability density function (Directionality PDF). Sample sizes, n = 8 except YAP^cKO^;TAZ^cKO^, n = 1

**Supplementary Figure 4.**
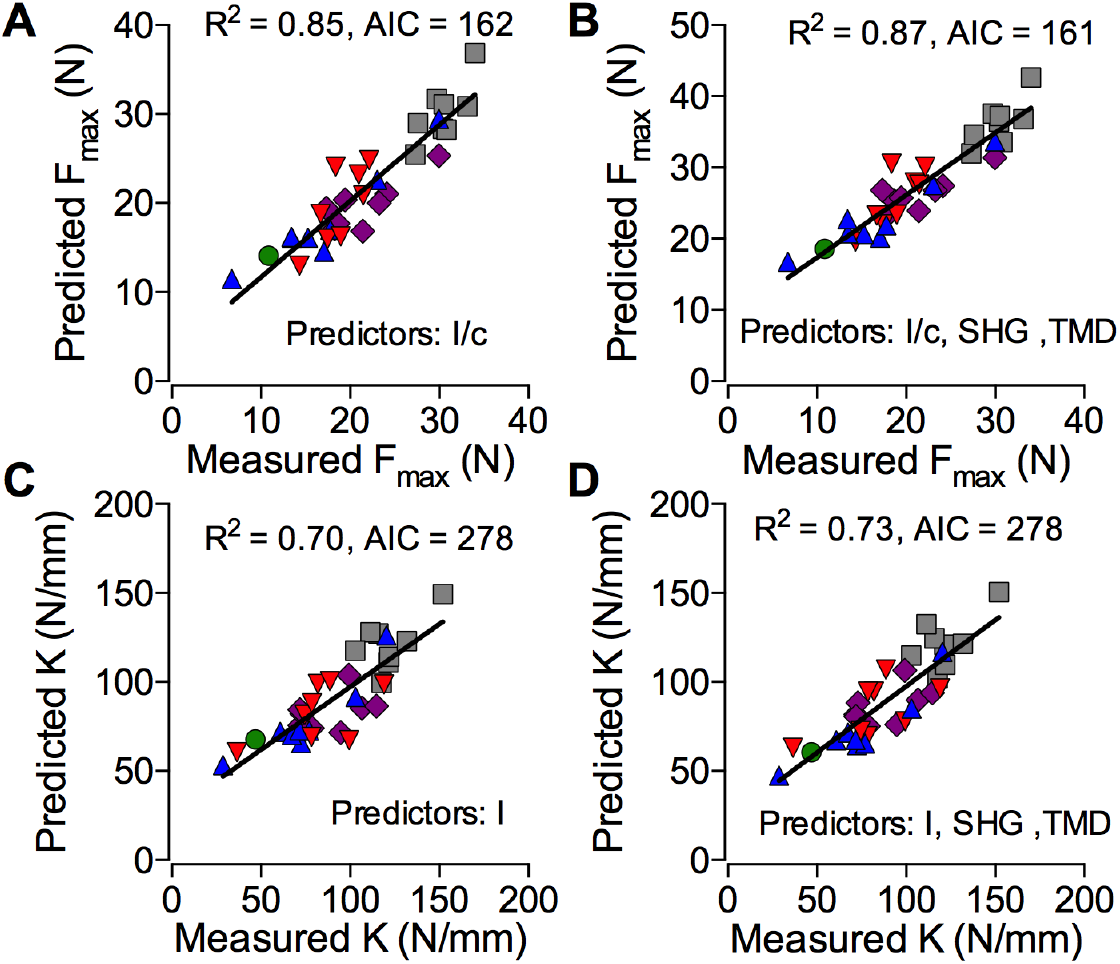
Best subsets analysis on morphological parameters from the YAP/TAZ ablation in Osterix-expressing cells from 8-week-old mice demonstrated increased goodness of fit parameters by accounting for collagen content and organization. Within the mice with Osterix driven deletion of YAP/TAZ, we compared experimental ultimate load (F_u_) with predicted model values with either (**A**) only section modulus (I/c) as a predictor or (**B**) the best model from the multivariate analysis, which incorporated both I/c and second harmonic generated (SHG) signal intensity. Similarly, we comparing experimental to predicted stiffness with either (**C**) only moment of inertia or (**D**) the best model from the multivariate analysis, which incorporated both moment of inertia and second harmonic generated signal intensity. Akaike’s information criterion (AIC) decreased while R^2^ increased in both regressions for maximum load and stiffness, suggesting a better goodness of fit with the addition of SHG. See methods for multivariate best subset analysis full details. Sample sizes, n = 8 except YAP^cKO^;TAZ^cKO^, n = 1

These data suggested that YAP and TAZ may regulate skeletal cell expression of collagen-related genes. To evaluate potential YAP/TAZ transcriptional targets, we first identified candidate genes whose mutations in humans cause osteogenesis imperfecta with similar penetrance to the observed mouse phenotypes (Table S2). For each candidate gene, we searched published ChIP-seq data in the UCSC Genome Browser^41^ for DNA binding domains of known YAP/TAZ co-transcription factors proximate (within 10 kb) to the transcription start site (TSS) (Table S2). We identified functional TEAD-binding MCAT elements (5’-CATTCCA/T-3’) in multiple OI-related genes. Based on the motif score^42^and TSS-proximity of each identified MCAT element, we selected Col1a1, Col1a2, and SerpinH1 as putative YAP/TAZ target genes. We performed quantitative PCR amplification of candidate mRNA transcripts isolated from 8 week-old femoral cortical bone preparations. Osterix-conditional YAP/TAZ deletion significantly reduced Col1a1 and SerpinH1 expression in a manner dependent on allele dosage (Fig. 5F,G). Differences in Col1a2 expression did not reach significance (Fig. 5H).

**Table S2:** UCSC Genome Browser candidate gene identification. Identification of functional TEAD-binding MCAT elements (5’-CATTCCA/T-3’) in multiple OI-related genes. *indicate statistically significant motif scores (p < 0.05). TSS = Transcriptional Start Site. + and − indicate downstream and upstream of TSS, respectively.

| Gene<br>(location) | OI-related gene<br>function<sup>43</sup> | TEAD binding<br>motif? | Motif<br>score<sup>42</sup> | Distance to<br>TSS (kb) |
| --- | --- | --- | --- | --- |
| *COL1A1*<br>(Chr17: 48,261,457) | Classical Sillence<br>types | Yes | 1.22 | +0.353 |
| *COL1A2*<br>(Chr7: 94,023,873) |  | Yes | 1.96*<br>1.72* | -1.303<br>-2.529 |
| *SERPINH1*<br>(Chr11: 75,271,275) | Collagen<br>chaperone defects | Yes | 1.58*<br>1.96* | +1.408<br>-2.570 |
| *FKBP10*<br>(Chr17: 39,966,270) |  | Yes | 1.27 | +8.071 |
| *CRTAP*<br>(Chr3: 43,212,006) | 3-Hydroxylation<br>defects | Yes | 1.27 | +8.071 |
| *LEPRE1*<br>(Chr1: 43,212,006) |  | Yes | 1.27 | +8.071 |
| *PPIB*<br>(Chr15: 64,448,014) |  | No |  |  |
| *OSTERIX/Sp7*<br>(Chr12: 53,717,808) | Osteoblast<br>maturation defects | No |  |  |

### YAP and TAZ are functionally redundant in DMP1-expressing cells and promote cortical and cancellous bone development

To determine the roles of YAP and TAZ late in the skeletal cell sequence, we next used 8kb-Dmp1-Cre ^33^ mice to preferentially delete YAP and TAZ from osteocytes^33,34^. We used the same breeding strategy as above to generate YAP/TAZ allele dosage-dependent DMP1-conditional knockouts. All pups were born at expected Mendelian ratios, and unlike the Osterix-conditional knockouts, double homozygous DMP1-cKO mice (cDKO^DMP1^) exhibited grossly normal skeletal structure and composition at P0 (Fig. 6A), without ribcage malformation (Fig. 6B) or spontaneous fractures. Femoral lengths, measured at 12 weeks of age (Fig.6C), revealed no effect of YAP/TAZ allele dosage on long bone growth, except for significantly reduced length in the double homozygous knockouts for both sexes (Fig.6D). A single copy of either gene was sufficient to rescue this defect, suggesting functional redundancy. Therefore, for further analyses, we selected for comparison littermate wild type (WT) and cDKO^DMP1^ mice. In metaphyseal cancellous bone of the distal femur, YAP/TAZ deletion from DMP1-expressing cells reduced trabecular number and increased trabecular spacing (Fig.6E-F). Femoral cortical bone exhibited reduced thickness and area, without changes in medullary area, indicating reduced periosteal bone in cDKO^DMP1^ mice (Fig.6G-H). Unlike Osterix-conditional knockouts, YAP/TAZ deletion from DMP1-expressing cells did not significantly alter cross sectional moment of inertia or tissue mineral density (Fig. 6H).

**Figure 6.**
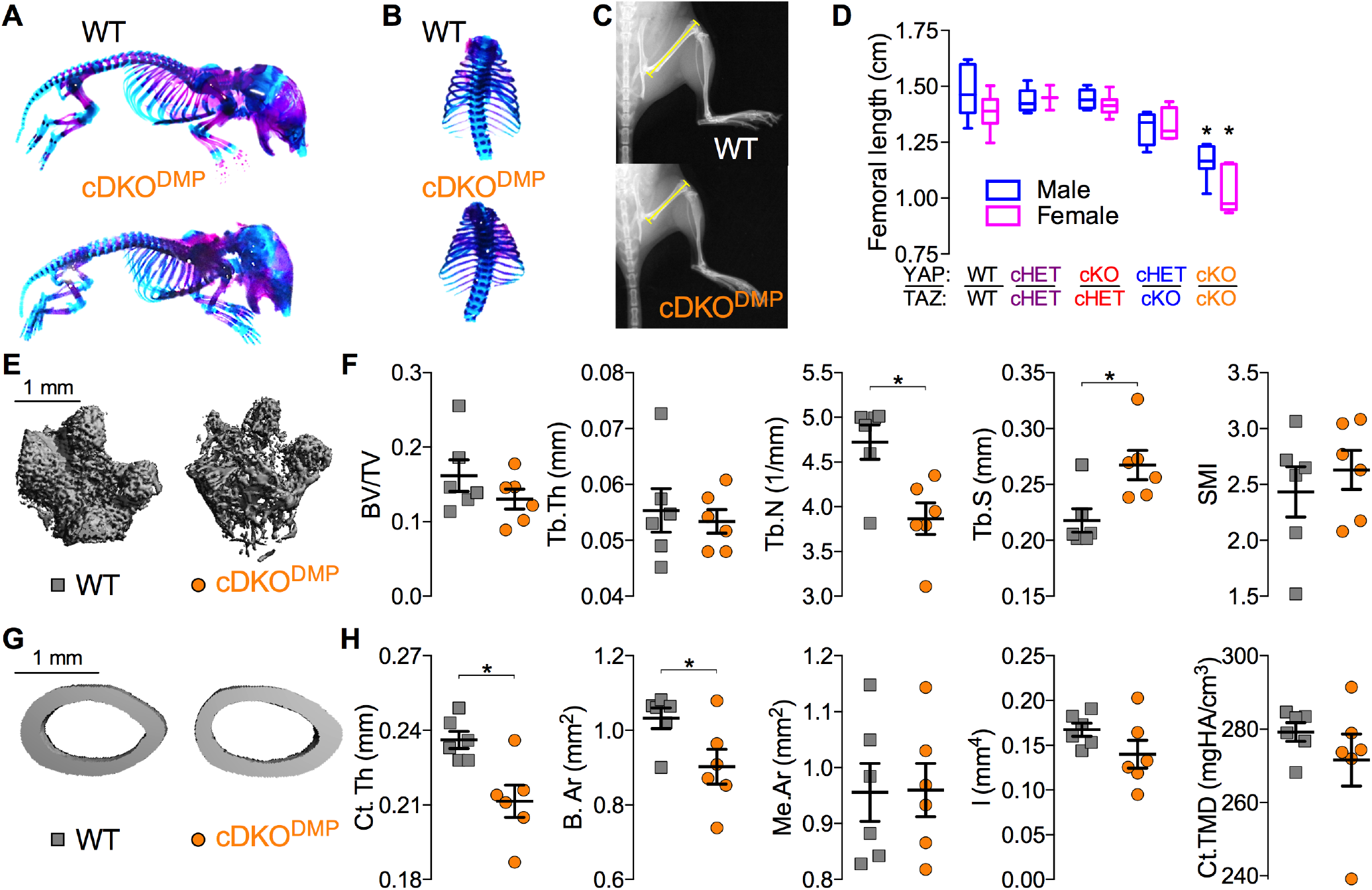
YAP/TAZ ablation from DMP1-expressing cells moderately impaired cancellous and cortical bone development. Littermate mice at P0 and P84 were selected for skeletal morphology analysis by skeletal preparation and microCT analysis. Whole body (**A**) and rib cage (**B**) skeletal preparations stained with Alcian blue/Alizarin red demonstrated normal skeletal structure in cDKO^DMP^ at P0. **C**) Representative radiographs illustrating femoral length at week 12. **D**) Femoral length was reduced only dual homozygous knockouts for both males and females. **E)** Representative microCT reconstructions of distal metaphyseal microarchitecture at week 12. **F)** YAP/TAZ deletion moderately altered cancellous bone microarchitecture: bone volume fraction (BV/TV), trabecular thickness (Tb.Th), number (Tb.N), spacing (Tb.Sp), and structural model index (SMI). **G)** Representative microCT reconstructions of femoral mid-diaphysis. **H)** YAP/TAZ deletion reduced cortical microarchitectural properties: bone area (B.Ar), medullary area (Me.Ar), cortical thickness (Ct.Th), moment of inertia in the direction of bending (I), and cortical tissue mineral density (Ct.TMD). Data are presented as individual samples with lines corresponding to the mean and standard error of the mean (SEM). Sample sizes, n = 6. Scale bars indicate 1 mm for microCT reconstructions.

### DMP1-conditional YAP/TAZ deletion impairs bone matrix collagen content, organization, and collagen-associated gene expression

Homozygous YAP/TAZ deletion from DMP1-expressing cells did not significantly reduce maximum load (Fig. 7A; p = 0.3) or bending stiffness (Fig. 7B; p = 0.08) in three-point bend testing to failure. Correlation of extrinsic properties with bone cross-sectional geometry (Fig. 7A,B) revealed significantly linear correlations for WT, but not for cDKO^DMP^ bones, though differences in regression lines between WT and cDKO^DMP^ did not reach statistical significance (p=0.4 and 0.1 for maximum load and stiffness, respectively). Matrix collagen content was significantly reduced in cDKO^DMP^ cortical bone while differences in the distribution of collagen fiber alignment did not reach statistical significance (p = 0.1) (Fig. 7C-F). To assess osteocyte density, the number osteocytes per bone area (Ot.N/B.Ar) were quantified by histomorphometry in cortical bone of the same limbs evaluated for microCT, mechanical testing, and SHIM. Cortical Ot.N/B.Ar was not significantly altered (p = 0.5), and differences in the percentage of empty lacunae were not significant (p = 0.1) (Fig. 7G-I). Best subsets regression analysis revealed significant contributions of both bone geometry (section modulus and moment of inertia) and collagen matrix properties (SHG) in predicting mechanical behavior (Fig. S5A-D).

**Figure 7.**
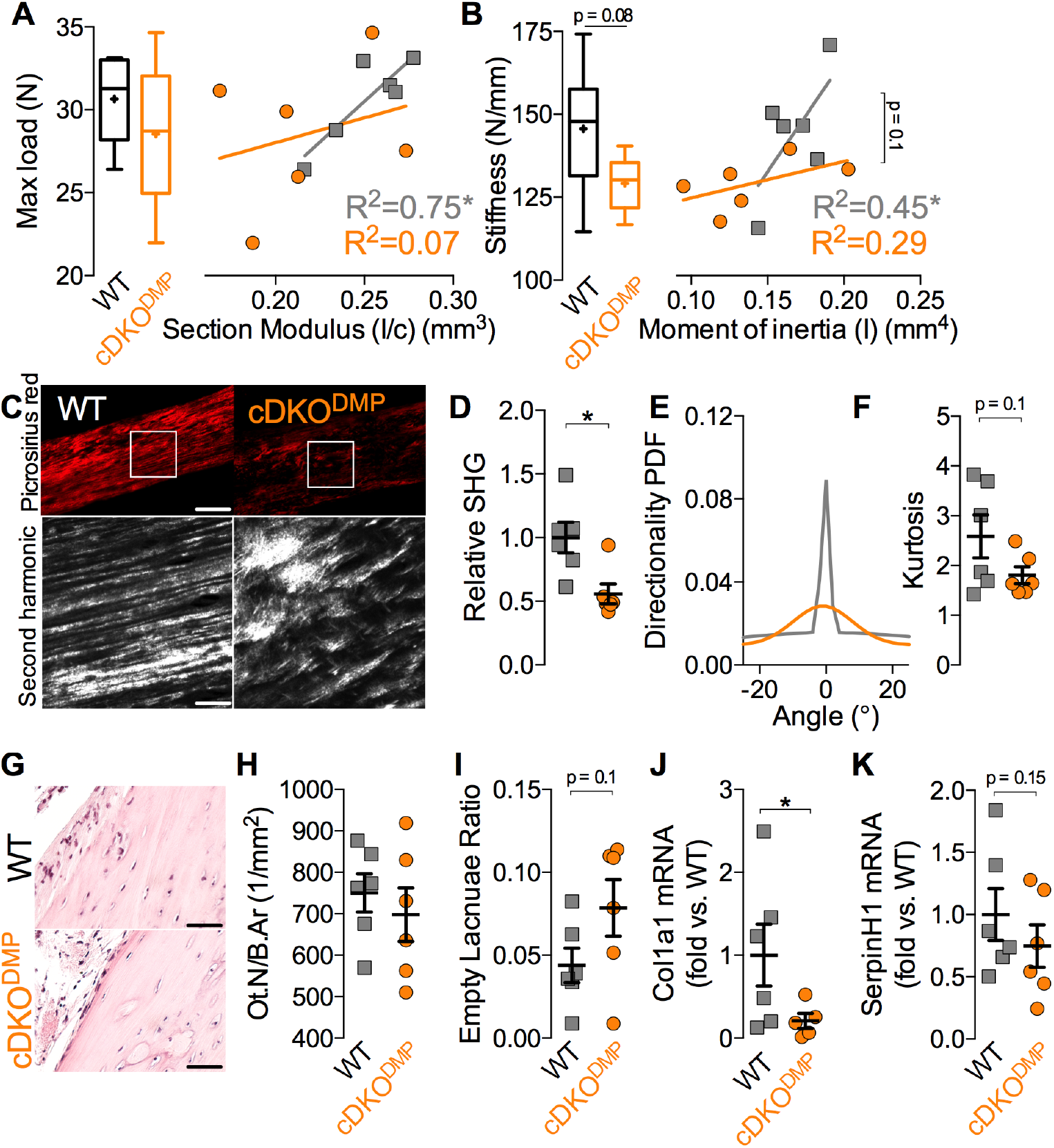
Dual homozygous YAP/TAZ deletion from DMP1-expressing cells reduced matrix collagen content, organization, and collagen-associated gene expression. Femora from 12 week-old littermate mice were evaluated by mechanical testing, second harmonic imaging microscopy, histomorphometry, and gene expression analyses. **A**) DMP1-conditional YAP/TAZ deletion did not significantly reduce maximum load or intrinsic failure properties. Reductions in extrinsic stiffness and intrinsic elastic properties did not reach significance (p = 0.08 and 0.1, respectively). Polarized light microscopy and second harmonic generation demonstrated reduced matrix collagen content and microstructural organization (**D**). The kurtosis (**E**) of the fiber directionality distributions (**F**) were not statistically significant (p = 0.01). Histological tissue sections of cortical bone stained with hematoxylin and eosin (H&E) (**G**) could not detect significant differences in osteocyte density expressed as osteocyte number per bone area (**H**) or the ratio of empty to total lacunae (**I**). YAP/TAZ deletion significantly reduced mRNA expression levels of Col1a1 (**J**), but SerpinH1 was not significant (p=0.15) (**K**). Sample sizes, n = 6 for all analyses. Scale bars equal 100, 25, and 10 microns in the Picrosirius red, SHG, and H&E images, respectively.

Like YAP/TAZ deletion from the full skeletal lineage, DMP1-conditional ablation reduced Col1a1 expression *in vivo* (Fig. 7J), with a similar trend for SerpinH1 (p = 0.15; Fig. 7K). Col1a2 was not significantly reduced (p = 0.8; data not shown). Unlike Osx-dependent knockouts, DMP1-mediated deletion also significantly reduced Runx2 expression (Fig. S6G).

**Supplementary Figure 5.**
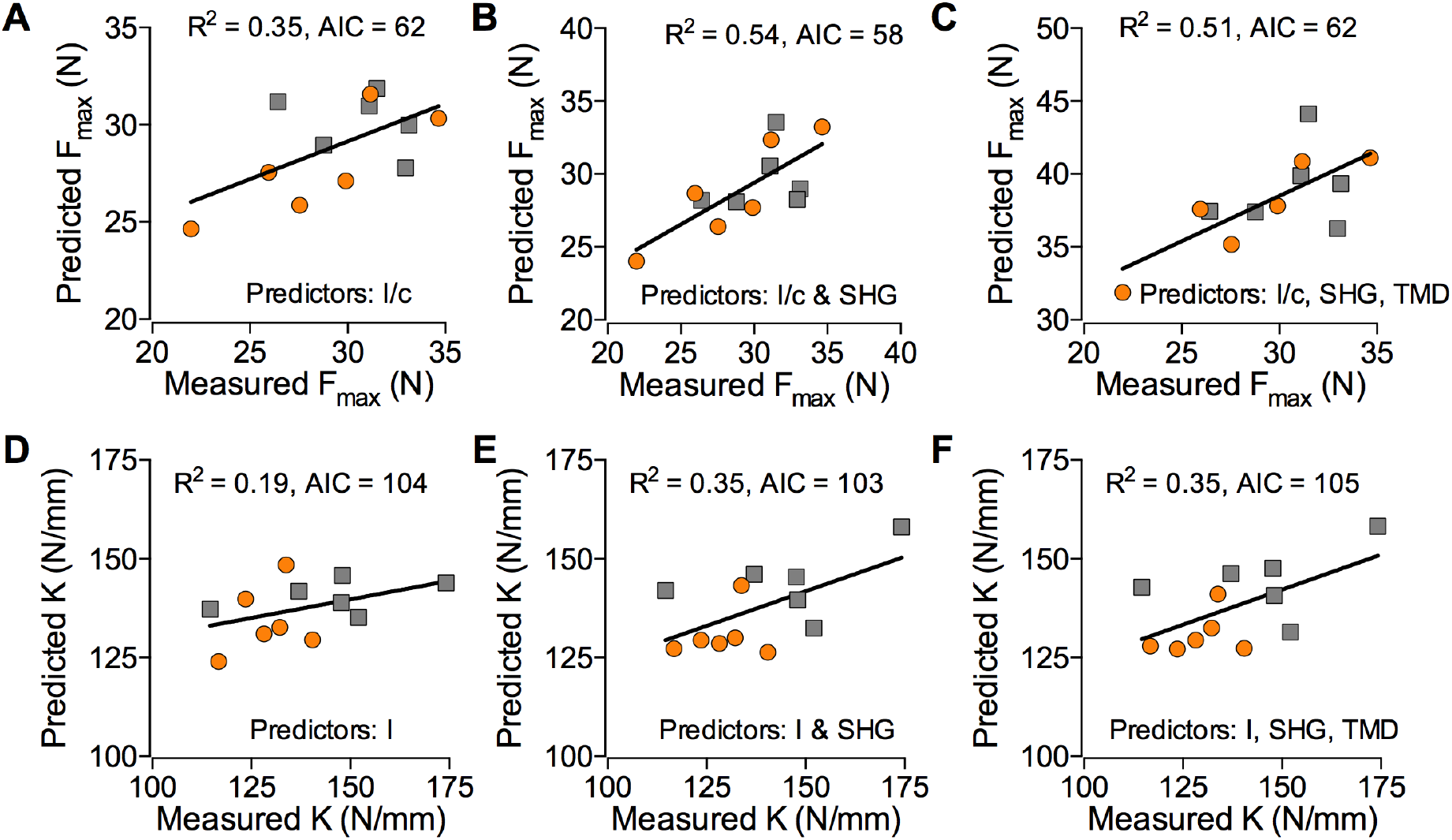
Best subsets analysis on morphological parameters from the YAP/TAZ ablation in DMP1-expressing cells from 12-week-old mice demonstrated enhanced goodness of fit parameters by accounting for collagen content and organization. Within the mice with DMP1-driven deletion of YAP/TAZ, we similarly compared experimental ultimate load (F_u_) with predicted model values with only moment of inertia as a predictor (**A**), moment of inertia and second harmonic generated signal intensity (**B**), or moment of inertia, second harmonic generated signal intensity, and tissue mineral density (**C**). Similarly, we compared experimental to predicted stiffness using the same predictors, respectively (**D-F**). Akaike’s information criterion (AIC) decreased while R^2^ increased in both regressions with maximum load and stiffness, suggesting optimal goodness of fit with the addition of SHG. See methods for full details of multivariate best subset analysis. Sample size, n = 6.

**Supplementary Figure 6.**
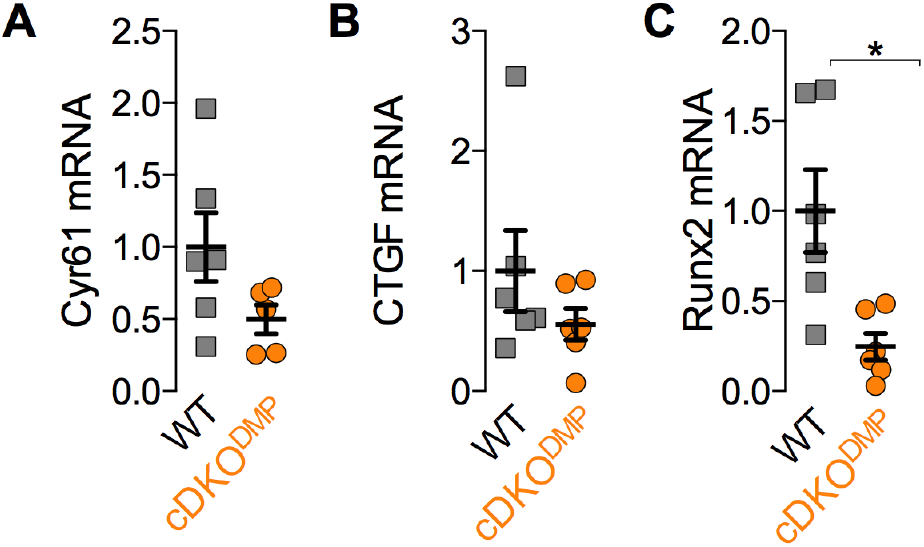
YAP/TAZ ablation in DMP1-expressing cells decreases trends in bone mRNA expression levels of canonical YAP/TAZ target genes. (**A**) CTGF, and (**B**) Cyr61 expression were down, yet not significant. **C**) Runx2 expression was only significantly down. mRNA was isolated from bone cell enriched lysates from femurs with DMP1-drive deletion of YAP/TAZ. Sample size, n = 6.

### Acute YAP/TAZ-TEAD inhibition reduced collagen and collagen-associated gene expression

Since changes in gene expression could result either from direct YAP/TAZ regulation or from developmental changes in bone cell populations, we next tested whether acute pharmacological YAP/TAZ inhibition using the small molecule verteporfin (VP) would also reduce YAP/TAZ activity and Col1a1 and SerpinH1 expression *in vitro* and *in vivo*. VP treatment of osteoblast-like UMR-106 cells *in vitro* reduced expression of the canonical YAP/TAZ-TEAD target gene, CTGF, and reduced synthetic YAP/TAZ-TEAD-responsive promoter activity (Fig. 8A)^26^. VP treatment *in vitro* dose-dependently reduced Col1a1 and Col1a2 expression, and SerpinH1 showed a similar but non-significant trend (p=0.1; Fig. 8B). WT mice were then treated every other day with intraperitoneal VP injection (100mg/kg) for two weeks. VP delivery significantly reduced expression of CTGF and SerpinH1 in liver tissue *in vivo*, but reductions in Col1a1 expression in liver and CTGF, Col1a1, and SerpinH1 in bone did not reach statistical significance (Fig. 8C,D). Consistent with genetic YAP/TAZ deletion, Col1a2 was not significantly reduced in either tissue.

**Figure 8.**
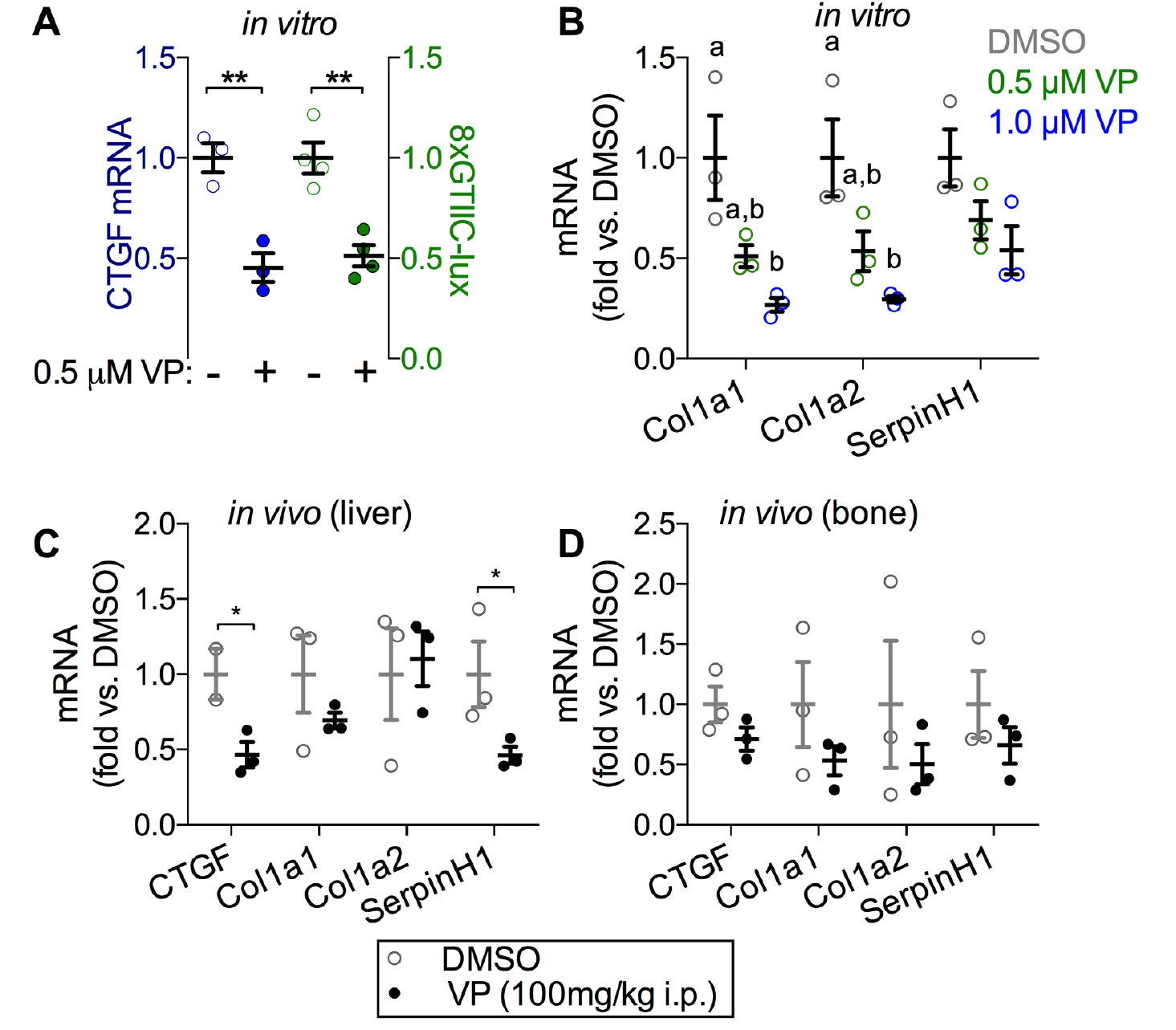
YAP/TAZ-TEAD inhibition with verteporfin reduced Col1a1 and SerpinH1 expression. **A**) Verteporfin (VP) treatment of UMR-106 cells reduced mRNA expression of the canonical YAP/TAZ-TEAD target gene *CTGF* commensurate with synthetic TEAD (8xGTIIC) reporter activity, demonstrating VP effectiveness. 8xGTIIC reporter activity was normalized to Renilla luciferase expression and expressed as fold vs. DMSO. (**B**) VP treatment dose dependently reduced mRNA levels of Col1a1, Col1a2, and SerpinH1 UMR-106 cells in comparison to DMSO. **C,D)** 16-week-old wild type mice received intraperinatal injections of VP (100 mg/kg) every two days. Livers and femora were harvested after 14 days to quantify mRNA expression. VP treatment significantly reduced hepatocyte expression of CTGF and SerpinH1 (**C**) and exhibited similar trends in bone (**D**), but differences were not statistically significant. Sample sizes, n = 3-4.

## Discussion

YAP and TAZ have been implicated as regulators of osteogenesis for over a decade^23,24^, and have been studied extensively *in vitro;* however, existing data on the differential roles of YAP vs. TAZ are strikingly contradictory^23–30,44–46^, and their combinatorial functions in bone have not been studied *in vivo*. To solve this apparent conflict in a physiological context, we used combinatorial skeletal cell-conditional YAP/TAZ deletion in mice to dissect the roles of YAP and TAZ both early and late in the skeletal cell differentiation sequence. Our data reveal that YAP and TAZ have mutually redundant, positive roles in both osteoblast progenitors and committed bone cells during development and regulate bone matrix mechanical properties putatively through transcriptional control of collagen expression and matrix organization.

### YAP/TAZ deletion from skeletal cells phenocopies osteogenesis imperfecta

YAP/TAZ deletion phenocopied osteogenesis imperfecta (OI), with severity dependent on targeted cell lineage and allele dosage. OI is a highly heterogeneous group of diseases characterized by bone fragility and deformity, whose severity varies from mildly increased fracture risk to perinatal lethality^47^. Though heritable through either autosomal dominant or recessive forms, mutations in the structure, organization, or quantity of collagen deposited in the bone matrix cause 85-90% of all OI cases^47,48^. The specific genetic mutations that cause the various types of OI remain poorly understood, though new putatively causal mutations continue to be discovered^49^. YAP/TAZ signaling has not been associated with OI, but elucidation of this pathway in bone may contribute new insights into the heterogeneity and/or etiology of the disease. Because global YAP deletion is embryonic lethal in animal models, loss-of-function mutations in YAP/TAZ are unlikely to cause human OI. However, there are many pathways that converge on YAP/TAZ which could place this signaling axis upstream of the human disease, including TGFβ-Smad2/3^11,12^ and WNT-β-catenin^9,10^. Further research will be required to evaluate whether YAP/TAZ signaling is causally linked to OI.

Mice possessing only a single copy of either gene in osterix-lineage cells rescued the lethality found in dual homozygous knockouts, demonstrating mutually compensatory function in skeletal cells. This contradicts some observations in the literature implicating divergent roles of YAP^23,26,27^ and TAZ^25,29,30^ in osteogenesis, but corroborate other reports of equivalent function^24,26,44,50^. These data also indicate that future loss-of-function studies aiming to understand the roles of these paralogs in bone must account for the activity of both genes in concert.

### YAP/TAZ regulate endochondral ossification

The spontaneous long bone fractures observed in Osterix-conditional YAP^cKO^;TAZ^cHET^ and YAP^cHET^;TAZ^cKO^ mice healed through natural endochondral repair similar to experimental neonatal fractures, which straighten and heal through bidirectional growth plate formation controlled by forces exerted by the adjacent muscle^51^. This suggests that this mechanotransductive realignment mechanism remained functional despite prior demonstration that YAP negatively regulates chondrogenesis both *in vitro*^52^ and *in vivo*^31^. However, the Osterix-Cre transgene is not strongly expressed in early or proliferating chondrocytes^32^, which are responsible for bone fragment re-alignment^51^. Instead, YAP/TAZ deletion from Osterix-expressing cells impaired endochondral ossification, potentially through reduced osteoprogenitor motility^26^, proliferation^52^, or survival^53^. Further research outside the scope of this study will be required to elucidate these mechanisms.

### YAP/TAZ regulate bone matrix collagen

Osterix-conditional knockout mice mimicked clinical observations of Types II and III OI and several established OI mouse models^43^, characterized by reduced bone volume, hypermineralization, and reduced intrinsic mechanical properties. For example, the human Col1a1 minigene mouse^54,55^, which expresses a human transgene with a clinically-observed mutation in pro-α_1_(I) collagen, dose-dependently reproduces the phenotypes seen here with Osterix1-conditional YAP/TAZ deletion. Like YAP^cKO^;TAZ^cKO^ mice, high dose pro-a_1_(I) transgenics exhibited neonatal lethality, while mice with moderate transgene dose mimicked the heterozygous knockouts with spontaneous femoral fractures and reduced failure, but not elastic bone material properties. Similarly, the naturally-occurring *oim* mouse, caused by a frameshift mutation in pro-a_2_(I) collagen, also features reduced bone mechanical properties and increased fracture incidence despite elevated mineral density^56,57^.

Notably, YAP/TAZ deletion later in the skeletal sequence (from DMP1-expressing cells) exhibited a similar but milder phenotype with moderate defects in cortical and cancellous bone accrual and matrix collagen content and organization. A single intact allele of either gene in DMP1-expressing cells sufficiently rescued the defect in long bone growth, indicating mutual YAP/TAZ redundancy.

### YAP/TAZ-dependent gene expression

Using a candidate-based approach selected from disease-causing mutations in human OI, we identified Col1a1 and SerpinH1 as YAP/TAZ-regulated genes in bone, with consistently reduced expression either by genetic YAP/TAZ ablation at multiple stages in the skeletal cell lineage or by acute YAP/TAZ-TEAD inhibition *in vitro* and *in vivo*. VP-induced reductions in CTGF, Col1a1, and SerpinH1 mRNA in bone did not reach significance, likely due to the small sample size and the less efficient biodistribution of porphryins to bone compared to liver and other tissues^58,59^. However, known YAP/TAZ target genes and SerpinH1 were significantly reduced in livers of VP-treated mice, indicating generality. These genes clearly do not constitute a sufficient set of all YAP/TAZ target genes with potential importance in bone development, and many other pathways are likely involved through other putative YAP/TAZ co-effectors^13–17^. Further research will be required to identify all YAP/TAZ-regulated genes in bone.

Both Osterix-Cre^60^ and DMP1-Cre^61^ transgenes have reports of targeting non-skeletal-lineage cell populations, which must be taken under consideration. However, the consistency in the phenotypes observed through *in vivo* YAP/TAZ ablation in DMP1-expressing cells and in Osterix-expressing cells, combined with the allele dosage-dependent response in bone phenotype severity, suggest the importance of skeletal cell YAP/TAZ. Taken together, these data suggest that YAP and TAZ redundantly promote bone matrix quality during development through transcriptional co-activation of TEAD to regulate Col1a1 and SerpinH1 expression *in vivo* and suggest that the YAP/TAZ signaling axis could have a role in osteogenesis imperfecta.

## MATERIALS AND METHODS

### Animals

Mice harboring loxP-flanked exon 3 alleles in both YAP and TAZ were kindly provided by Dr. Eric Olson (University of Texas Southwestern Medical Center). The tetracycline responsive B6.Cg-Tg(Sp/7-tTA,tetO-EGFP/cre)1AMc/J (Jackson laboratories) mice (Osx-Cre) were raised, bred, and evaluated without tetracycline administration to induce constitutive gene recombination in osteoprogenitor cells and their progeny^32^. Mature osteoblasts and osteocytes were separately targeted using Cre-recombination under the control of an 8kb fragment of the Dmp1 promoter (Dmp1-Cre)^33^. Mice with homozygous floxed alleles for both YAP and TAZ (YAP^fl/fl^;TAZ^fl/fl^) were mated with double heterozygous conditional knockout mice (YAP^fl/^+;TAZ^fl/^+;Osx-Cre and YAP^fl/+^;TAZ^fl/+^;Dmp1-Cre) to produce eight possible genotypes in each litter (Table S1). Both male and female mice were evaluated. No differences in skeletal phenotype were observed based on sex (Fig. S2A,B). All mice were fed regular chow ad libitum and housed in cages containing 2-5 animals each. Mice were maintained at constant 25°C on a 12-hour light/dark cycle. Mice were tail or ear clipped after weaning or prior to euthanasia and genotyped by PCR or using a third party service (Transnetyx Inc.) All protocols were approved by the Institutional Animal Care and Use Committee at the University of Notre Dame in adherence to federal guidelines for animal care.

### Skeletal preparations

Skeletal preparations stained with Alizarin red and Alcian blue were performed as described previously^62^. Briefly, mice were euthanized via CO_2_ asphyxiation at postnatal day 0 (P0) and skin, viscera, and adipose tissues were dissected quickly. Dissected skeletons were fixed in 80% ethanol and dehydrated in 95% ethanol overnight at room temperature immediately followed by incubation in 100% acetone for two days to remove remaining adipose tissue. Prepared skeletons were stained at room temperature using Alcian blue (A3157; Sigma) and Alizarin red (A5533; Sigma) for two days. Stained skeletons were incubated in 95% ethanol for 1 hour then remaining tissue was cleared through incubation in a 1% potassium hydroxide (KOH) solution. Skeletal preparations were stabilized in a 1% KOH / 20% glycerol solution and imaged in 1% KOH / 50% glycerol solution with a Nikon D7100 camera.

### μCT and mechanical testing

Femora were harvested at 8 or 12 weeks of age and stored at -20°C until evaluation. The frozen specimens were thawed and imaged in PBS using a vivaCT 80 scanner (Scanco Medical) to determine trabecular and cortical femoral bone architecture prior to mechanical testing to failure in three-point bending. The mid-diaphysis and distal femur were imaged with an X-ray intensity of 114 μA, energy of 70 kVp, integration time of 300 ms, and resolution of 10 μm. Mid-diaphyseal and distal femoral 2D tomograms were manually contoured, stacked and binarized by applying a Gaussian filter (sigma =1, support = 1) at a threshold value of 250 mg HA/cm^3^. Mechanical testing analysis of the femurs was carried out by three-point bend testing. The femurs were loaded with the condyles facing down onto the bending fixtures with a lower span length of 4.4 mm. The upper fixture was aligned with the mid-diaphysis. The femora were loaded to failure at a rate of 0.5 mm/s using the ElectroForce 3220 Series testing system (TA Instruments). Sample sizes: n = 8 animals for each group except n = 1 animal for the YAP^cKO^;TAZ^cKO^.

### Histology

Bone samples were fixed in 10% neutral buffered formalin overnight at 4°C, decalcified in a 7:3 mixture of 14% neutral buffered EDTA solution and 10% neutral buffered formalin for 3 weeks at 4°C, and transferred to 70% ethanol for storage at 4°C. Paraffin sections of 5 μm thickness for both histological staining and immunohistochemistry were deparaffinized and rehydrated in a graded series of ethanol solutions and washed three times in PBS. For immunohistochemistry, heat induced antigen retrieval in a 1x sodium citrate buffer preceded the quenching of endogenous peroxides with PerxoBlock (Invitrogen). Primary antibodies were compared to negative control sections. Anti-Osterix (abcam: Cat# ab22552, 1:250), anti-YAP (Cell Signaling: Cat# 14074, 1:400), anti-TAZ (Cell Signaling: Cat# 4883, 1:400) were both applied overnight at 4°C and compared to sections receiving only secondary antibody. Colorimetric detection using the DAB Peroxidase HRP-linked Substrate Kit (Vector Labs) allowed for immunohistochemical detection of YAP, TAZ, or Osterix-positive cells. Hematoxalyin and eosin stains (H&E) were used to detect nuclei and tissue morphology, Safranin O/Fast Green (Saf-O) to visualize sulfated glycosaminogylcans, and Picrosirius Red to visualize collagen fibers. Osteocyte number per bone area and empty lacanuae ratio (empty lacunae over both empty and full lacnaue were manually counted using ImageJ (NIH) on H&E stained sections. Hypertrophic chondrocyte zone percent thickness (HZ thickness %) was calculated manually in ImageJ (NIH). Three separate lines were drawn across the area of positive Saf-O staining, normalized to the respective length of the hypertrophic chondrocyte zone within each line, and averaged together for each image. Three images per animal were taken at the growth plate. Sample sizes: n = 8 animals for each group except n = 1 animal for the YAP^cKO^;TAZ^cKO^.

### Imaging

Histological and immunohistochemical sections were imaged on a 90i Upright/Widefield Research Microscope (Nikon Instruments) at the 4x, 10x, 20x, and 40x objectives. Three-point bend femur sections stained with Picro-sirius Red were imaged under polarized light using an Eclipse ME600 Microscope (Nikon Instruments) at the 20x objective while second harmonic image microscopy (SHIM) images were taken on a multiphoton-enabled Fluoview Research Microscope (Olympus) at 875nm with the 25x objective on sections oriented in the same direction for all groups. All SHIM images were quantified using ImageJ (NIH) and reported as mean pixel intensity within the cortical region of the femoral bone. Mean pixel intensities across four separate regions of interest within each image of the cortex were averaged as technical replicates for a given histological section. Directionality was assessed by using the FIJI macro to ImageJ (NIH). Raw directionality histograms were imported from FIJI to MATLAB (v2015.b). The histograms from each image were averaged and then fit with a Gaussian function assuming a normal distribution. The generated average probability density functions were reported for each genotype.

### Verteporfin delivery in vivo

Six littermate control (four male and two female) mice (YAP^WT^;TAZ^WT^) were aged until 16 weeks. Two males and one female were then either administered Verteporfin (100 mg/kg) in 0.9% saline or dimethyl sulfoxide (DMSO) in 0.9% saline by intraperinatal injection every other for day for 2 weeks. Livers and femurs from both verteporfin-treated and DMSO-treated mice were harvested on the day of the last injection.

### RNA isolation and qPCR

Total bone and liver samples were snap frozen in liquid nitrogen cooled isopentane for 1 minute before quickly being store at -80°C until processed. Upon processing, tissue was homogenized via mortar and pestle and RNA from the sample was collected using Trizol Reagant (Life Technologies Inc.) followed with centrifugation in chloroform. Upon layer separation, the RNA sample was purified using the RNA Easy Kit (Qiagen) and quantified by spectrophotometry using a NanoDrop 2000. Reverse transcriptase polymerase chain reaction (RT-PCR) was performed on 0.5 ug/ul concentration of RNA using the TaqMan Reverse Transcription Kit (Thermo Fisher Scientific). Quantitative polymerase chain reaction (qPCR) assessed mRNA amount using a CFX Connect (Bio-Rad) relative to the internal control of glyceraldehyde 3-phosphate dehydriogenase (GAPDH). Data are presented using the ΔΔCt method. Specific mouse primer sequences are listed in Table S2. n = 6 animals for each group except n = 1 animal for the YAP^cKO^;TAZ^cKO^.

**Table S3:**
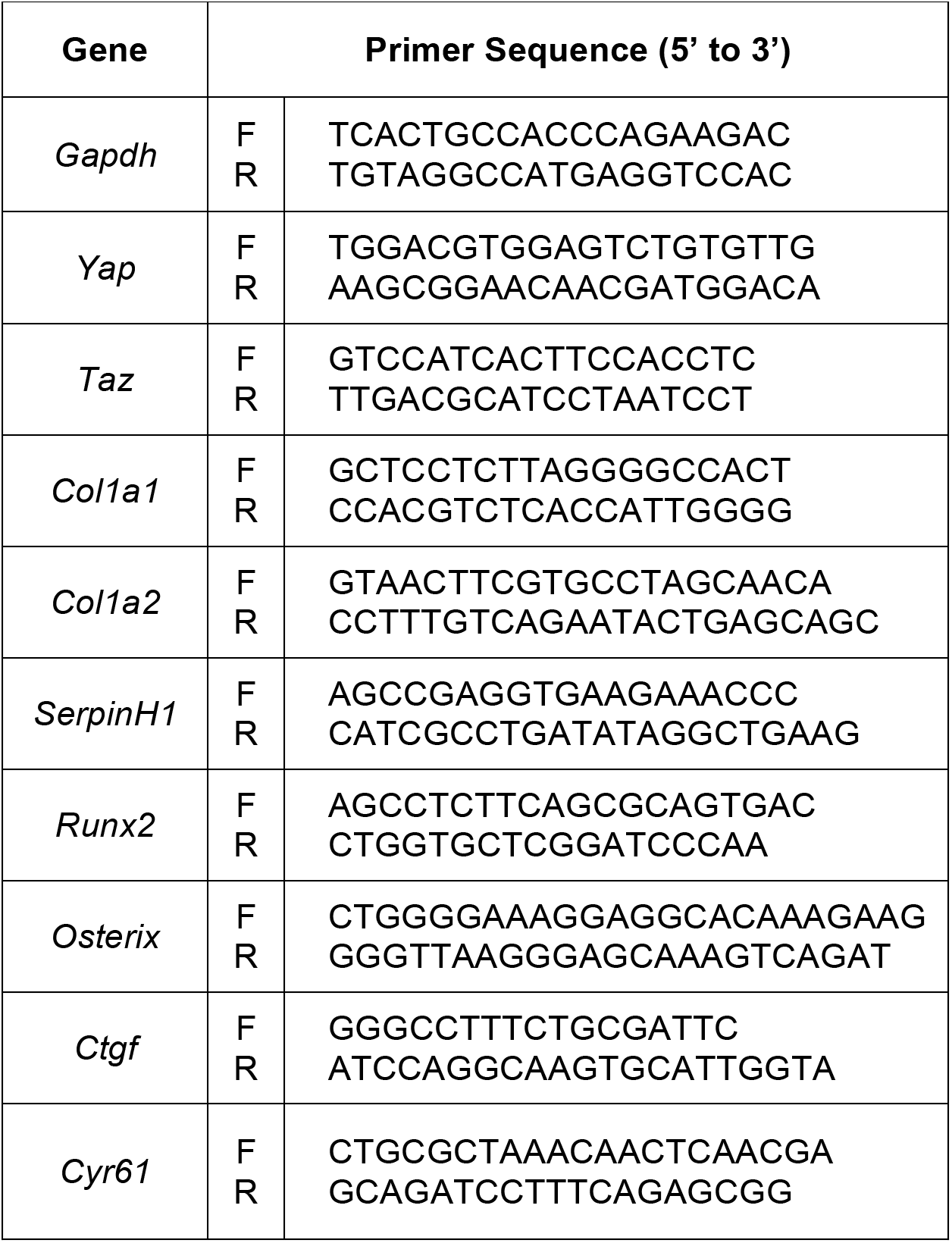
Mouse primer sequences for qPCR.

### Rat osteosarcoma cell line

UMR-106 cells were cultured in Dulbecco’s Modified Eagle’s Medium (DMEM) modified to contain 4 mM L-glutamine, 4500 mg/L glucose, 1 mM sodium pyruvate, and 1500 mg/L sodium bicarbonate and 10% FBS according to the American Type Culture Collection (ATCC; Catalog #: 30-2002) recommendations. UMR-106 cells at 50 % confluence were transfected with the TEAD-responsive luciferase reporter plasmid 8XGTIIC-Luciferase (Addgene) and a control Renilla plasmid, both kindly provided by Munir Tanas, MD, University of Iowa. Forty-eight hours after transfection, cells were treated with either DMSO, 0.5 μM, or 1μM verteporfin in serum-free conditions for 1 hour. All verteporfin experiments were carried out in the dark. Cells were then lysed immediately using the Dual-Luciferase Reporter Assay System according to manufactures instructions (Promega). Luciferase activity was measured on a VICTOR 3 (Perkin Elmer) plate reader and normalized to baseline renilla activity as previously described^63^. Separately cultured UMR-106 cells were simultaneously treated with either DMSO, 0.5 μM, or 1μM verteporfin and then cultured under serum-free conditions for one hour. mRNAs were then isolated and purified with the RNA-easy Mini-Kit (Qiagen) following the manufacturer’s instructions. mRNA (0.5 μg) was reverse transcribed using the TaqMan Reverse Transcription Kit (Thermo Fisher Scientific). Quantitative polymerase chain reaction (qPCR) assessed mRNA amount using a CFX Connect (Bio-Rad) relative to the internal control of glyceraldehyde 3-phosphate dehydriogenase (GAPDH). Data are presented using the ΔΔCt method. Specific rat primer sequences are listed in Table S3. Sample size, n = 3 independent experiments.

**Table S4:**
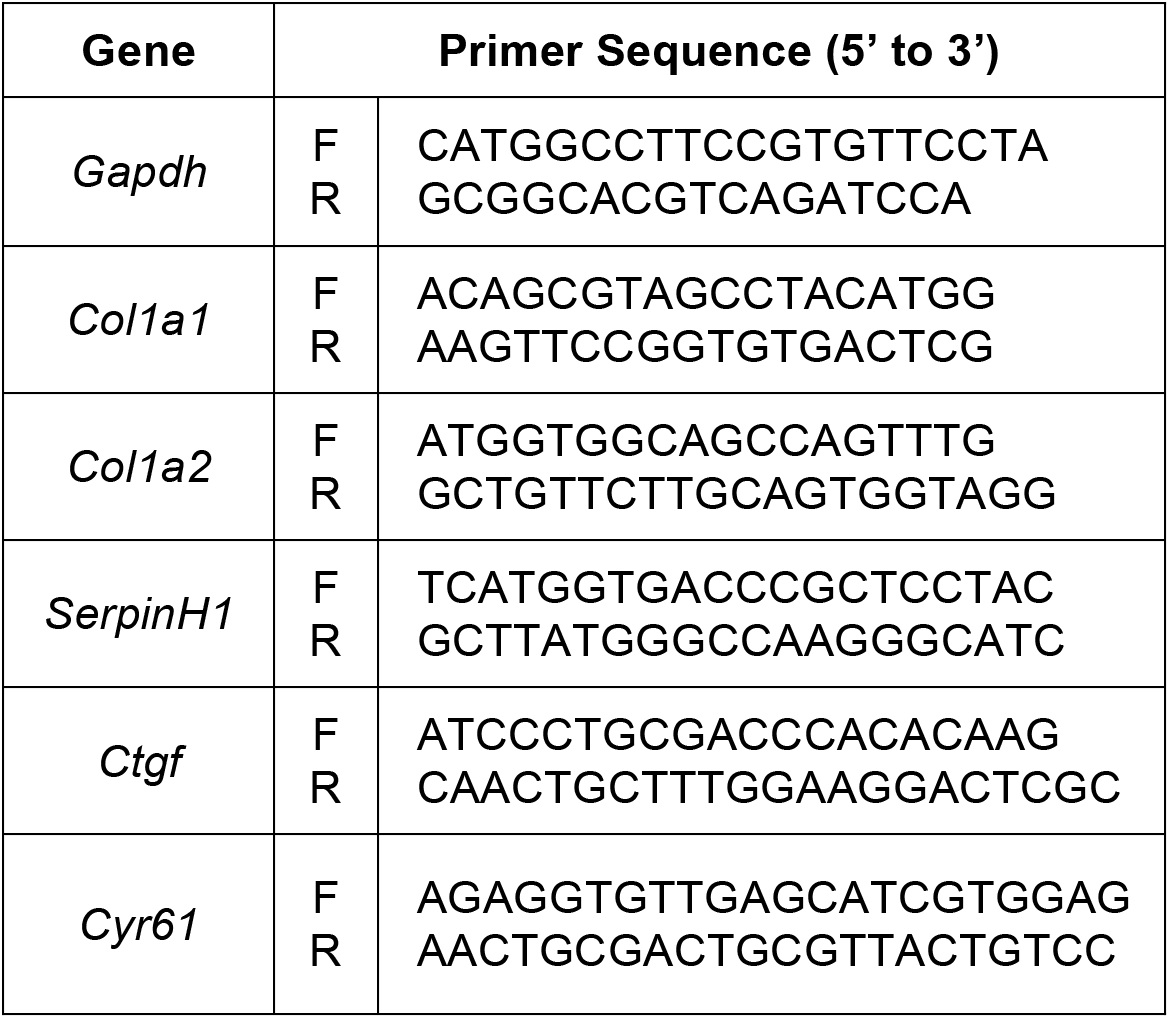
Rat primer sequences for qPCR.

### Statistics and regression

All statistics and linear regressions including the ANCOVA were performed in GraphPad Prism. Comparisons between two groups were made using the independent t-test while comparisons between 3 or more groups were made using a one-way ANOVA with post-hoc Bonferroni’s multiple comparisons test if the data were normally distributed according to D’Agostino-Pearson omnibus normality test and homoscedastic according to Bartlett’s test. When parametric test assumptions were not met, data were log-transformed prior and residuals were evaluated. If necessary, the non-parametric Kruskal-Wallis test with post-hoc Dunn’s multiple comparisons was used. A p-value of less than 0.05 (adjusted for multiple comparisons) was considered significant. Data are represented as individual samples with mean ± standard error of the mean (s.e.m.). Multivariate correlations were perfomed using the *bestglm* package (available at https://cran.r-project.org/web/packages/bestglm/ version 0.36) in the open-source GNU statistical package R (Version 2.13.1). We followed a modified procedure previously described ^40^. Briefly, we chose to use an exhaustive best subsets algorithm to determine the best predictors of maximum load and stiffness from a subset of morphological parameters measured, which included moment of inertia (I) or section modulus (I/c), tissue mineral density (TMD), and second harmonic generation (SHG) intensity based on the Akaike’s information criterion (AIC).^39^ The lowest AIC selects the best model while giving preference to less complex models (those with fewer explanatory parameters). Finally, the overall “best” model for each predicted mechanical property was compared to the prediction from only the moment of inertia (I/c or I for maximum load and stiffness, respectively) using Type II general linear regression.

## References

1. de Crombrugghe, B., Lefebvre, V. & Nakashima, K. Regulatory mechanisms in the pathways of cartilage and bone formation. Curr. Opin. Cell Biol. 13, 721–7 (2001).

2. Karsenty, G. Transcriptional Control of Skeletogenesis. Annu. Rev. Genomics Hum. Genet. 9, 183–196 (2008).

3. Javed, A., Chen, H. & Ghori, F. Y. Genetic and transcriptional control of bone formation. Oral Maxillofac. Surg. Clin. North Am. 22, 283–93, v (2010).

4. Varelas, X. The Hippo pathway effectors TAZ and YAP in development, homeostasis and disease. Development 141, 1614–1626 (2014).

5. Vassilev, A., Kaneko, K. J., Shu, H., Zhao, Y. & DePamphilis, M. L. TEAD/TEF transcription factors utilize the activation domain of YAP65, a Src/Yes-associated protein localized in the cytoplasm. Genes Dev. 15, 1229–41 (2001).

6. Zhao, B. et al. TEAD mediates YAP-dependent gene induction and growth control. Genes Dev. 22, 1962–71 (2008).

7. Yagi, R., Chen, L.-F., Shigesada, K., Murakami, Y. & Ito, Y. A WW domain-containing Yes-associated protein (YAP) is a novel transcriptional co-activator. EMBO J. 18, 2551–2562 (1999).

8. Rosenbluh, J. et al. β-Catenin-driven cancers require a YAP1 transcriptional complex for survival and tumorigenesis. Cell 151, 1457–1473 (2012).

9. Heallen, T. et al. Hippo pathway inhibits Wnt signaling to restrain cardiomyocyte proliferation and heart size. Science 332, 458–61 (2011).

10. Azzolin, L. et al. YAP/TAZ Incorporation in the β-Catenin Destruction Complex Orchestrates the Wnt Response. Cell 158, 157–170 (2014).

11. Varelas, X. et al. TAZ controls Smad nucleocytoplasmic shuttling and regulates human embryonic stem-cell self-renewal. Nat. Cell Biol. 10, 837–848 (2008).

12. Varelas, X. et al. The Crumbs complex couples cell density sensing to Hippo-dependent control of the TGF-β-SMAD pathway. Dev. Cell 19, 831–44 (2010).

13. Komori, T. et al. Targeted Disruption of Cbfa1 Results in a Complete Lack of Bone Formation owing to Maturational Arrest of Osteoblasts. Cell 89, 755–764 (1997).

14. Otto, F. et al. Cbfa1, a Candidate Gene for Cleidocranial Dysplasia Syndrome, Is Essential for Osteoblast Differentiation and Bone Development. Cell 89, 765–771 (1997).

15. Mundlos, S. et al. Mutations Involving the Transcription Factor CBFA1 Cause Cleidocranial Dysplasia. Cell 89, 773–779 (1997).

16. Kato, M. et al. Cbfa1-independent decrease in osteoblast proliferation, osteopenia, and persistent embryonic eye vascularization in mice deficient in Lrp5, a Wnt coreceptor. J. Cell Biol. 157, (2002).

17. Afzal, F. et al. Smad function and intranuclear targeting share a Runx2 motif required for osteogenic lineage induction and BMP2 responsive transcription. J. Cell. Physiol. 204, 63–72 (2005).

18. Hong, W. & Guan, K.-L. The YAP and TAZ transcription co-activators: Key downstream effectors of the mammalian Hippo pathway. Semin. Cell Dev. Biol. 23, 785–793 (2012).

19. Morin-Kensicki, E. M. et al. Defects in yolk sac vasculogenesis, chorioallantoic fusion, and embryonic axis elongation in mice with targeted disruption of Yap65. Mol. Cell. Biol. 26, 77–87 (2006).

20. Hossain, Z. et al. Glomerulocystic kidney disease in mice with a targeted inactivation of Wwtr1. Proc. Natl. Acad. Sci. 104, 1631–1636 (2007).

21. Miesfeld, J. B. et al. Yap and Taz regulate retinal pigment epithelial cell fate. Development 142, (2015).

22. Xin, M. et al. Hippo pathway effector Yap promotes cardiac regeneration. Proc. Natl. Acad. Sci. U. S. A. 110, 13839–44 (2013).

23. Zaidi, S. K. et al. Tyrosine phosphorylation controls Runx2-mediated subnuclear targeting of YAP to repress transcription. EMBO J. 23, 790–9 (2004).

24. Hong, J.-H. et al. TAZ, a Transcriptional Modulator of Mesenchymal Stem Cell Differentiation. Science (80-.). 309, (2005).

25. Hong, J.-H. & Yaffe, M. B. TAZ: A β-Catenin-like Molecule that Regulates Mesenchymal Stem Cell Differentiation. Cell Cycle 5, 176–179 (2006).

26. Dupont, S. et al. Role of YAP/TAZ in mechanotransduction. Nature 474, 179–83 (2011).

27. Seo, E. et al. SOX2 Regulates YAP1 to Maintain Stemness and Determine Cell Fate in the Osteo-Adipo Lineage. Cell Rep. 3, 2075–2087 (2013).

28. Park, H. W. et al. Alternative Wnt Signaling Activates YAP/TAZ. Cell 162, 780–794 (2015).

29. Byun, M. R. et al. Canonical Wnt signalling activates TAZ through PP1A during osteogenic differentiation. Cell Death Differ. 21, 854–863 (2014).

30. Yang, J.-Y. et al. Osteoblast-Targeted Overexpression of TAZ Increases Bone Mass In Vivo. PLoS One 8, e56585 (2013).

31. Deng, Y. et al. Yap1 Regulates Multiple Steps of Chondrocyte Differentiation during Skeletal Development and Bone Repair. Cell Rep. 14, 2224–2237 (2016).

32. Rodda, S. J. & McMahon, A. P. Distinct roles for Hedgehog and canonical Wnt signaling in specification, differentiation and maintenance of osteoblast progenitors. Development 133, 3231–3244 (2006).

33. Bivi, N. et al. Cell autonomous requirement of connexin 43 for osteocyte survival: Consequences for endocortical resorption and periosteal bone formation. J. Bone Miner. Res. 27, 374–389 (2012).

34. Delgado-Calle, J. et al. Control of Bone Anabolism in Response to Mechanical Loading and PTH by Distinct Mechanisms Downstream of the PTH Receptor. J. Bone Miner. Res. 32, 522–535 (2017).

35. Jepsen, K. J., Silva, M. J., Vashishth, D., Guo, X. E. & van der Meulen, M. C. H. Establishing biomechanical mechanisms in mouse models: practical guidelines for systematically evaluating phenotypic changes in the diaphyses of long bones. J. Bone Miner. Res. 30, 951–66 (2015).

36. Kourtis, L. C., Carter, D. R. & Beaupre, G. S. Improving the Estimate of the Effective Elastic Modulus Derived from Three-Point Bending Tests of Long Bones. Ann. Biomed. Eng. 42, 1773–1780 (2014).

37. Guss, J. D. et al. Alterations to the Gut Microbiome Impair Bone Strength and Tissue Material Properties. J. Bone Miner. Res. (2017). doi:10.1002/jbmr.3114

38. Chen, X., Nadiarynkh, O., Plotnikov, S. & Campagnola, P. J. Second harmonic generation microscopy for quantitative analysis of collagen fibrillar structure. Nat. Protoc. 7, 654–669 (2012).

39. Akaike, H. A new look at the statistical model identification. IEEE Trans. Automat. Contr. 19, 716–723 (1974).

40. Schneider, P., Voide, R., Stampanoni, M., Donahue, L. R. & Müller, R. The importance of the intracortical canal network for murine bone mechanics. Bone 53, 120–128 (2013).

41. Kent, W. J. et al. The human genome browser at UCSC. Genome Res. 12, 996–1006 (2002).

42. Daily, K., Patel, V. R., Rigor, P., Xie, X. & Baldi, P. MotifMap: integrative genome-wide maps of regulatory motif sites for model species. BMC Bioinformatics 12, 495 (2011).

43. Forlino, A., Cabral, W. A., Barnes, A. M. & Marini, J. C. New perspectives on osteogenesis imperfecta. Nat. Rev. Endocrinol. 7, 540–557 (2011).

44. Tang, Y. et al. MT1-MMP-Dependent Control of Skeletal Stem Cell Commitment via a β1-Integrin/YAP/TAZ Signaling Axis. Dev. Cell 25, 402–416 (2013).

45. Kim, K. M. et al. Shear Stress Induced by an Interstitial Level of Slow Flow Increases the Osteogenic Differentiation of Mesenchymal Stem Cells through TAZ Activation. PLoS One 9, e92427 (2014).

46. Kim, M., Kim, T., Johnson, R. L. & Lim, D.-S. Transcriptional co-repressor function of the hippo pathway transducers YAP and TAZ. Cell Rep. 11, 270–82 (2015).

47. Forlino, A. & Marini, J. C. Osteogenesis imperfecta. Lancet 387, 1657–1671 (2016).

48. Rauch, F. & Glorieux, F. H. Osteogenesis imperfecta. Lancet 363, 1377–1385 (2004).

49. Ackermann, A. M. & Levine, M. A. Compound heterozygous mutations in *COL1A1* associated with an atypical form of type I osteogenesis imperfecta. Am. J. Med. Genet. Part A (2017). doi:10.1002/ajmg.a.38238

50. Zhong, W. et al. Mesenchymal Stem Cell and Chondrocyte Fates in a Multishear Microdevice Are Regulated by Yes-Associated Protein. Stem Cells Dev. 22, 2083–2093 (2013).

51. Rot, C., Stern, T., Blecher, R., Friesem, B. & Zelzer, E. A Mechanical Jack-like Mechanism Drives Spontaneous Fracture Healing in Neonatal Mice. Dev. Cell 31, 159–170 (2014).

52. Karystinou, A. et al. Yes-associated protein (YAP) is a negative regulator of chondrogenesis in mesenchymal stem cells. Arthritis Res. Ther. 17, 147 (2015).

53. Halder, G., Dupont, S. & Piccolo, S. Transduction of mechanical and cytoskeletal cues by YAP and TAZ. Nat. Rev. Mol. Cell Biol. 13, 591–600 (2012).

54. Khillan, J. S., Olsen, A. S., Kontusaari, S., Sokolov, B. & Prockop, D. J. Transgenic mice that express a mini-gene version of the human gene for type I procollagen (COL1A1) develop a phenotype resembling a lethal form of osteogenesis imperfecta. J. Biol. Chem. 266, 23373–9 (1991).

55. Pereira, R., Khillan, J. S., Helminen, H. J., Hume, E. L. & Prockop, D. J. Transgenic mice expressing a partially deleted gene for type I procollagen (COL1A1). A breeding line with a phenotype of spontaneous fractures and decreased bone collagen and mineral. J. Clin. Invest. 91, 709–16 (1993).

56. Chipman, S. D. et al. Defective pro alpha 2(I) collagen synthesis in a recessive mutation in mice: a model of human osteogenesis imperfecta. Proc. Natl. Acad. Sci. U. S. A. 90, 1701–5 (1993).

57. Vanleene, M. et al. Ultra-structural defects cause low bone matrix stiffness despite high mineralization in osteogenesis imperfecta mice. Bone 50, 1317–1323 (2012).

58. Richter, A. M., Cerruti-Sola, S., Sternberg, E. D., Dolphin, D. & Levy, J. G. Biodistribution of tritiated benzoporphyrin derivative (3H-BPD-MA), a new potent photosensitizer, in normal and tumor-bearing mice. J. Photochem. Photobiol. B. 5, 231–44 (1990).

59. Akens, M. K. et al. Photodynamic Therapy of Vertebral Metastases: Evaluating Tumor-to-Neural Tissue Uptake of BPD-MA and ALA-PpIX in a Murine Model of Metastatic Human Breast Carcinoma. Photochem. Photobiol. 83, 1034–1039 (2007).

60. Chen, J. et al. Osx-Cre Targets Multiple Cell Types besides Osteoblast Lineage in Postnatal Mice. PLoS One 9, e85161 (2014).

61. Zhang, J. & Link, D. C. Targeting of Mesenchymal Stromal Cells by *Cre*-Recombinase Transgenes Commonly Used to Target Osteoblast Lineage Cells. J. Bone Miner. Res. 31, 2001–2007 (2016).

62. McLeod, M. J. Differential staining of cartilage and bone in whole mouse fetuses by alcian blue and alizarin red S. Teratology 22, 299–301 (1980).

63. Tanas, M. R. et al. Mechanism of action of a WWTR1(TAZ)-CAMTA1 fusion oncoprotein. Oncogene 35, 929–38 (2016).

